# Fate and plasticity of SARS-CoV-2-specific B cells during memory and recall response in humans

**DOI:** 10.1101/2022.10.07.511336

**Authors:** Yves Zurbuchen, Jan Michler, Patrick Taeschler, Sarah Adamo, Carlo Cervia, Miro E. Raeber, Ilhan E. Acar, Jakob Nilsson, Michael B. Soyka, Andreas E. Moor, Onur Boyman

## Abstract

B cell responses to different pathogens recruit tailored effector mechanisms, resulting in functionally specialized subsets. For human memory B cells (MBCs), these include CD21^+^ resting, CD21^−^CD27^+^ activated, and CD21^−^CD27^−^ atypical cells. Whether these subsets follow deterministic or interconnected fates is unknown. We demonstrate in COVID-19 patients that single clones of SARS-CoV-2-specific MBCs followed multiple fates with distinctive phenotypic and functional characteristics. 6–12 months after infection, most circulating MBCs were CD21^+^ resting cells, which also accumulated in peripheral lymphoid organs where they acquired markers of tissue residency. Conversely, at acute infection and following SARS-CoV-2-specific immunization, CD21^−^ MBCs became the predominant subsets, with atypical MBCs expressing high T-bet, inhibitory molecules, and distinct chemokine receptors. B cell receptor sequencing allowed tracking of individual MBC clones differentiating into CD21^+^, CD21^−^CD27^+^, and CD21^−^CD27^−^ cell fates. Collectively, single MBC clones can adopt functionally different trajectories, thus contributing to immunity to infection.

## INTRODUCTION

Upon encounter with cognate antigens, B and T lymphocytes are endowed with the capacity to form memory cells^1, 2^. Memory lymphocytes are usually long lived and provide faster and more vigorous immune responses upon secondary contact with their specific antigen^3^. Some memory cells circulate between blood, secondary lymphoid organs and bone marrow, while others migrate to peripheral tissues and mucosal sites where they can become tissue resident^4^. Whereas subdivision of labor in terms of tissue homing and effector functions has been well characterized for memory T cells, functionally different subsets also exist in memory B cells (MBCs). Antigen-stimulated B cells receiving instructive signals from their interaction with CD4^+^ T helper (Th) cells can further differentiate in germinal centers (GCs) of secondary lymphoid organs or via an extrafollicular pathway. In the GC, this differentiation includes affinity maturation through somatic hypermutation (SHM) of the B cell receptor (BCR), following which B cells can become long-lived plasma cells or MBCs^5–7^. Most long-lived plasma cells home to bone marrow niches where they are able to continuously secrete high-affinity antibodies protective against a homologous pathogen^8, 9^, whereas resting GC-derived MBCs encode a broader repertoire and are subsequently able to provide protection against variants of the initial pathogen^10,11^. Upon reencounter with their cognate antigen, MBCs differentiate into antibody-secreting plasma cells or reenter GCs where they undergo additional _SHM_^12, 13^.

MBCs can be subdivided into phenotypically and functionally distinct subsets^14^. In humans, resting MBCs typically express high surface levels of CD21, also known as complement receptor 2, and express the tumor necrosis factor (TNF) receptor superfamily member CD27. Contrarily, absence of CD21 expression marks CD21^−^CD27^+^ activated and CD21^−^CD27^−^ ‘atypical’ MBCs, both of which represent class-switched B cell subsets^15–17^. Unlike resting MBCs, the origin and differentiation path of activated and, particularly, atypical MBCs is less well defined. CD21^−^CD27^+^ B cells are thought to represent a GC-derived population prone to plasma cell differentiation^18^. Conversely, CD21^−^CD27^−^ atypical MBCs have been found in chronic infection, immunodeficiency, and autoimmune diseases where they are thought to be of extrafollicular origin^19–25^. CD21^−^CD27^+^ activated and CD21^−^CD27^−^ atypical antigen-specific MBCs have been detected transiently after different vaccines^15, 16, 18, 26, 27^ and during infections with certain pathogens^26, 28–30^, including acute severe acute respiratory syndrome coronavirus 2 (SARS-CoV-2)^31–35^. Atypical MBCs are characterized by expression of the transcription factor T-bet, which is essential for their development, as well as high abundance of CD11c and several inhibitory coreceptors, such as Fc receptor-like (FcRL) protein 5 (FcRL5)^36–38^. Atypical MBCs have been shown to be able to differentiate into antibody-secreting cells^29, 39^. Thus, atypical MBCs could play a role in protective immune responses.

Here, we studied antigen-specific MBC subsets in human subjects at different time points after infection with SARS-CoV-2 and in individuals following SARS-CoV-2-specific vaccination. We found SARS-CoV-2 spike-binding (spike^+^) CD21^−^CD27^+^ activated MBCs were the predominant subset in circulation during acute infection and upon vaccination, with substantial contribution of atypical MBCs, whereas at 6–12 months after infection CD21^+^ resting MBCs became prevalent. By using single-cell RNA sequencing (scRNA-seq), we discovered that single B cell clones were able to adopt different MBC subset fates and functional signatures upon antigen reexposure.

## RESULTS

### Longitudinal kinetics of SARS-CoV-2-specific MBC responses

To study antigen-specific MBC subsets in a human setting of natural infection and controlled immunization, we recruited a longitudinal cohort of coronavirus disease 2019 (COVID-19) patients at acute infection and at six and 12 months after infection (referred to as memory phase) (Fig. 1a). Of the 65 patients, 42 had mild COVID-19 and 23 severe COVID-19. Thirty-five patients received a SARS-CoV-2 mRNA vaccination between the six- and 12-month time point, and three subjects were vaccinated between acute infection and the six-month time point (Supp. Table 1). We developed a multimer staining based on biotinylated spike and receptor-binding domain (RBD) proteins of SARS-CoV-2 to analyze antigen-specific MBCs with a 28-color spectral flow cytometry panel (Fig. 1b, Supp. Fig. 1a). Analyzing the non-vaccinated samples, we observed a strong increase in frequencies of spike^+^ and RBD^+^ MBCs following SARS-CoV-2 infection, which remained stably high up to one year after infection (Fig. 1c–e). Frequencies of spike^+^ MBCs were comparable in mild and severe COVID-19 patients (Fig. 1f).

**Fig. 1.**
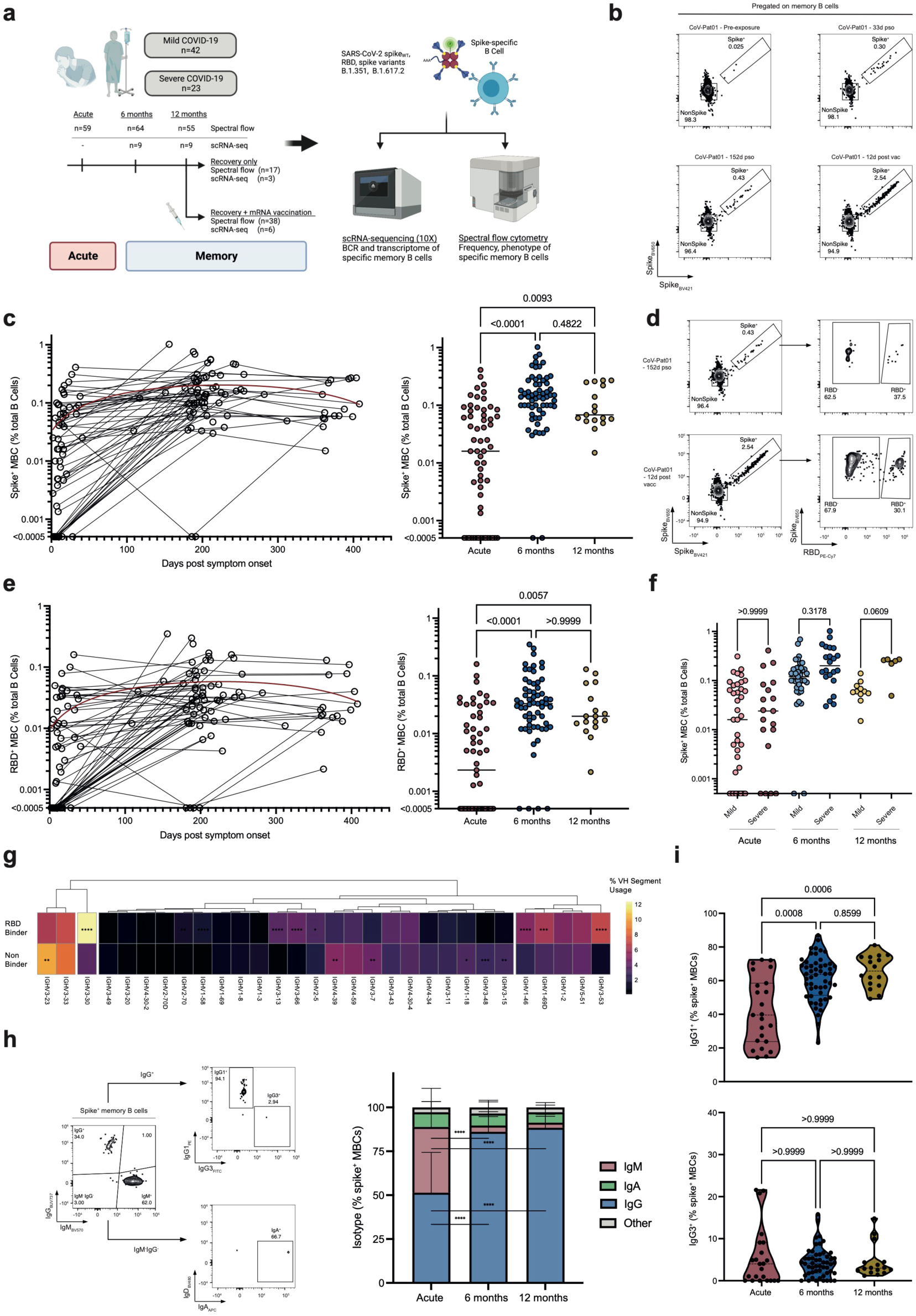
Longitudinal analysis of kinetics and magnitude of antigen-specific memory B cells upon SARS-CoV-2 infection. **a**, Overview of study design and cohort. **b**, Representative flow cytometry plots of spike multimer-stained memory B cells (MBCs; gating strategy shown in Supp. Fig. 1a) of the same individual (CoV-Pat01) shown before SARS-CoV-2 infection (top left), at 33 d (top right) and 152 d (bottom left) post symptom onset (pso), and at 12 d after vaccination (bottom right). Numbers indicate percentages of parent population. **c**, Frequency of spike^+^ MBCs at indicated time points pso (left). Paired samples are connected with lines. Second-order polynomial function (red line, R^2^=0.1932) is fitted to the data. Dot plots and medians (right) of frequencies of spike^+^ B cells at acute infection (n=59) and six (n=61) and 12 months (n=17) after infection. Samples collected after vaccination were excluded from this analysis. **d**, Representative flow cytometry plots showing gating strategy for RBD^+^ and spike^+^ MBCs of the same individual as in **b**. Numbers indicate percentages of parent population. **e**, Frequency of RBD^+^ MBCs at indicated time points pso (left). Paired samples are connected with lines. Second-order polynomial function (red line, R^2^=0.1298) is fitted to the data. Dot plots and medians (right) of frequencies of RBD^+^ B cells at acute infection (n=59) and six (n=61) and 12 months (n=17) after infection. **f**, Frequency of spike^+^ B cells at acute infection and six and 12 months after infection, separated by disease severity, comparing mild (acute n=40, six months n=39, 12 months n=11) and severe COVID-19 (acute n=19, six months n=22, 12 months n=6). **g**, Heatmap comparing V heavy (VH) gene usage between RBD binders and non-binders. VH are sorted by hierarchical clustering, with colors indicating frequencies. The 30 most frequently used segments in RBD binders are shown. **h**, Representative gating strategy of indicated isotypes in spike^+^ MBCs of a patient at acute infection (left). Stacked bar plot and mean + SD (right) showing isotype of spike^+^ MBCs at acute infection (n=23) and six (n=52) and 12 months (n=16) after infection. For all phenotypical analysis shown, samples were included if >9 spike^+^ MBCs were available. **i**, Violin plots of percentages of IgG1^+^ (top) and IgG3^+^ (bottom) spike^+^ MBCs at acute infection (n=23) and six (n=52) and 12 months (n=16) after infection. Samples were compared using a Kruskal-Wallis test with Dunn’s multiple comparison (**c**, **e**, **f**, **h**, **i**). Adjusted p-values are shown. In **g** frequencies were compared using two-proportions z-test with Bonferroni-based multiple testing correction. In **g** and **h** p-values are shown if significant (p<0.05). *p<0.05, **p<0.01, ***p<0.001, **** p<0.0001.

We purified spike^+^ versus spike^−^ MBCs by fluorescence-activated cell sorting from blood of nine patients at the memory phases (Supp. Fig. 1b, Supp. Table 2), followed by droplet-based scRNA-seq combined with feature barcoding and BCR sequencing. MBCs specific for RBD, wild-type spike (spike_WT_) or the spike variants B.1.351 (beta) and B.1.617.2 (delta) were identified by streptavidin multimers carrying oligonucleotide barcodes. The vast majority of spike variant^+^ and RBD^+^ MBCs also recognized spike_WT_ (Supp. Fig. 2a). Furthermore, we observed comparable frequencies of RBD-binding MBCs within spike^+^ MBCs using our sequencing approach as with flow cytometry (Supp. Fig. 2b,c). When analyzing V heavy and light chain frequencies of RBD^+^ MBCs, we found several chains, including *IGHV3-30*, *IGHV3-53*, *IGHV3-66*, *IGKV1-5, IGKV1-9* and *IGKV1-33*, to be enriched compared to RBD^−^ MBCs (Fig. 1g, Supp. Fig. 2d), which have been described to encode for RBD-binding antibodies^40–42^.

Moreover, we characterized immunoglobulin (Ig) isotypes and subtypes of spike^+^ MBCs by flow cytometry. During acute infection spike^+^ MBCs mainly expressed IgM and IgG, whereas IgG^+^ MBCs predominated the memory phases, mostly of the IgG1 subtype, and around 5–10% expressed IgA (Fig. 1h,i). In summary, these data identify a durable, spike-specific, and IgG1-dominated MBC response upon SARS-CoV-2 infection.

### Evolution of spike^+^ MBC subsets in blood from acute to memory phases

To further characterize the SARS-CoV-2-specific MBC response, we visualized spike^+^ MBCs by uniform manifold approximation and projection (UMAP) plots and performed an unsupervised Phenograph clustering (Fig. 2a, Supp. Fig. 3a–c). The UMAP grouped MBCs into IgM^+^, IgG^+^, and IgA^+^ cells (Fig. 2a). Also, it revealed a phenotypical shift from acute infection to the memory time points, which was driven by increasing CD21 expression, whereas T-bet, Blimp-1, CD11c, CD71, and FcRL5 expression diminished (Fig. 2b, Supp. Fig. 3b,c). The unsupervised Phenograph analysis of spike^+^ MBCs identified distinct clusters, such as the CD21^−^CD27^−^ atypical (MC07) cluster, which was Tbet^hi^, CD11c^+^ and FcRL5^+^, and the CD21^−^CD27^+^ activated (MC03) cluster characterized by a high expression of CD71, Blimp-1 and Ki-67 (Supp. Fig. 3a–c).

**Fig. 2.**
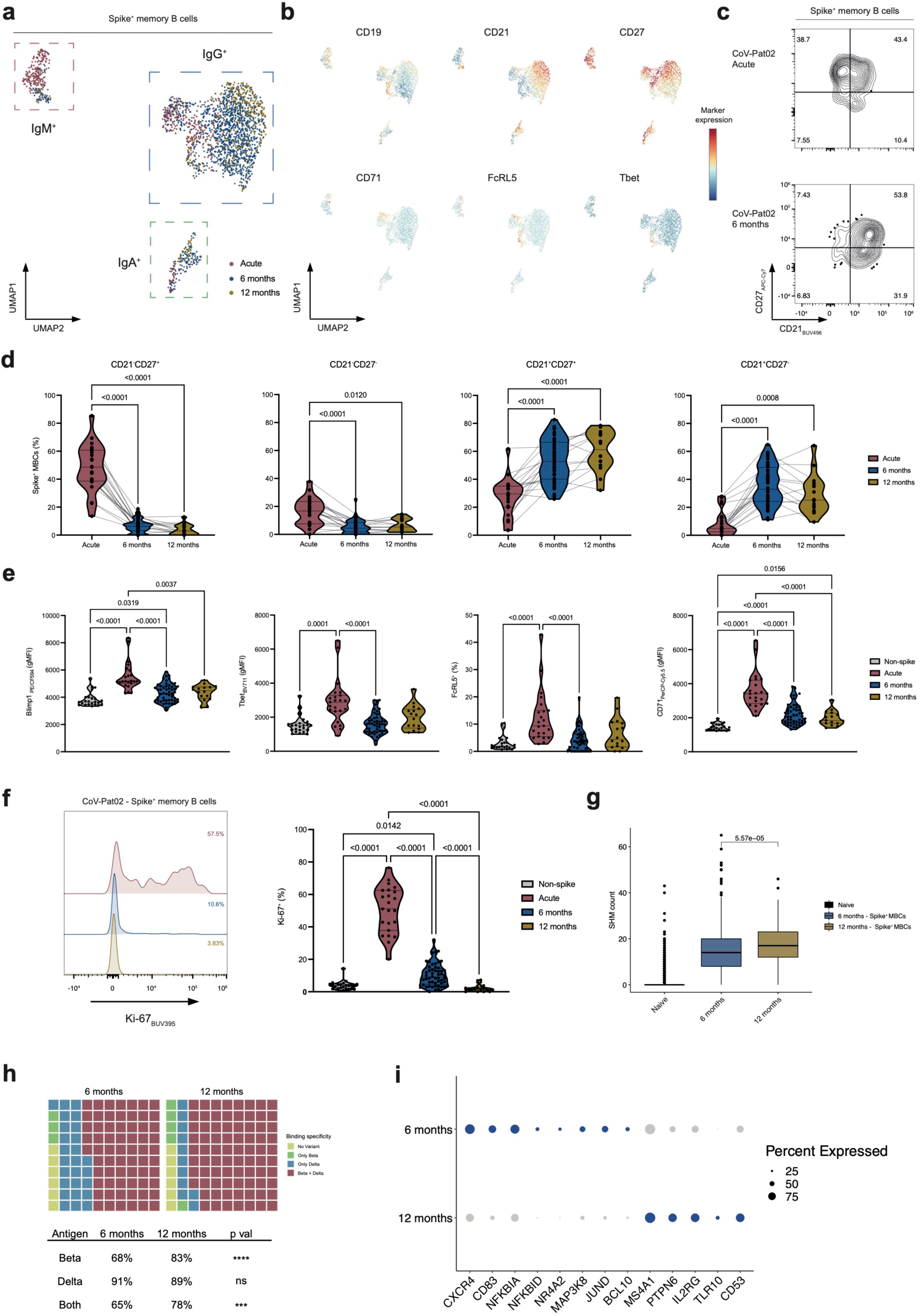
Phenotypic and functional characterization of circulating SARS-CoV-2-specific MBCs at acute and memory time points. **a**, Uniform manifold approximation and projection (UMAP) plots of spike^+^ MBCs (n=120, excluding samples after vaccination), subsampled to maximally 25 cells per sample and colored by time point. Islands of IgM^+^, IgA^+^ and IgG^+^ MBCs are indicated. **b**, As in **a** colored by indicated marker expression. **c**, Flow cytometry plots of spike^+^ MBCs during acute infection and six months after infection of the same patient (CoV-Pat02). **d**, Violin plots of frequencies of indicated subsets of spike^+^ MBCs at acute infection (n=23) and six (n=52) and 12 months (n=16) after infection. Paired samples are connected with lines. **e**, Violin plots of geometric mean fluorescence intensities (gMFI) or percentages of indicated markers in spike^+^ MBCs at acute infection (n=23) and six (n=52) and 12 months (n=16) after infection, compared to spike^−^ MBCs at acute infection (n=23). **f**, Representative histograms of Ki-67 in CoV-Pat02 (left) and violin plots of percentages of Ki-67^+^ spike^+^ MBCs (right) at indicated time points and compared to spike^−^ MBCs. **g**, Somatic hypermutation (SHM) counts of all samples (excluding those after vaccination), comparing SHM counts in spike^+^ MBCs that bound to any spike construct including wild-type (WT), variant, and RBD, at six (n=9) and 12 months (n=3) after infection, with naïve B cells serving as reference. **h**, Waffle plot of spike ^+^ MBCs binding beta (B.1.351) and delta spike variant (B.1.617.2) in unvaccinated individuals (n=9 at six and n=3 at 12 months). **i**, Expression (blue indicates upregulated genes) and percentages of selected, differently expressed genes in spike^+^ MBCs between six and 12 months. Dot size indicates frequency of positive cells. Samples in **d**–**f** were compared using a Kruskal-Wallis test with Dunn’s multiple comparison correction. Adjusted p-values are shown if significant (p<0.05). In **g**, two-sided Wilcoxon test was used with Holm multiple comparison correction. The box plots show median; box limits, interquartile range (IQR); whiskers, 1.5xIQR and outliers. In **h**, samples were compared using two-proportions z-test and, in **i**, using Wilcoxon Rank Sum test with Bonferroni correction. In **i**, all genes had adj. p<0.05 for differential expression between the two groups. ns p>0.05, *p<0.05, **p<0.01, ***p<0.001, **** p<0.0001.

These changes in CD21 and CD27 could be reproduced by manual gating at acute infection, six and 12 months thereafter (Fig. 2c). Spike^+^ CD21^−^CD27^+^ and, to a lesser extent, also CD21^−^CD27^−^ B cells were predominant during the acute response to SARS-CoV-2, but they were strongly reduced at six and 12 months after infection (Fig. 2d). Conversely, at the memory phases, CD21^+^CD27^+^ and CD21^+^CD27^−^ made up most of the antigen-specific MBC compartment (Fig. 2d). These dynamics were similar in patients with mild and severe COVID-19, except that severe COVID-19 patients had slightly higher levels of spike^+^ CD21^−^CD27^−^ B cells at six months after infection (Supp. Fig. 3d).

The transcription factors Blimp-1 and T-bet as well as FcRL5 and the activation marker CD71 were increased on spike^+^ B cells during acute infection and decreased at the memory phases (Fig. 2e). These changes were paralleled by strong proliferation of spike^+^ B cells during the acute phase, as indicated by high Ki-67 expression (Fig. 2f). Intriguingly, spike^+^ B cells continued to show lower but still significantly increased proliferation at six months after infection, which only returned to steady-state background levels at 12 months after infection (Fig. 2f).

Furthermore, we found a significantly increased SHM count in spike^+^ MBCs at 12 compared to six months in our scRNA-seq dataset (Fig. 2g). This difference was paralleled by an improved binding breadth measured by variant-binding capabilities of spike ^+^ MBCs (Fig. 2h). On a transcriptional level, MBCs at six months had upregulated genes associated with B cell activation and GC emigration ^43^, such as *NKFBIA*, *NFKBID*, *JUNB*, *MAP3K8* and *CD83*, compared to 12 months where *TLR10* and *IL2RG* were upregulated (Fig. 2i). Collectively, these data showed that SARS-CoV-2 infection induced a stable, resting CD21^+^ MBC population in the circulation, which continuously matured after infection.

### Circulating versus tonsillar SARS-CoV-2-specific MBC subsets

Having observed the dynamics of MBC subsets in blood, we wanted to assess the changes of SARS-CoV-2-specific B cells in a peripheral lymphoid organ. To this end, we obtained paired tonsil and blood samples of individuals only immunized with a SARS-CoV-2 mRNA vaccine but not infected with SARS-CoV-2 and of subjects that had recovered from SARS-CoV-2 infection and some of which were also vaccinated (Supp. Fig. 4a,b, Supp. Table 3). Using our multimer probe approach (Supp. Fig. 4a), we observed spike^+^ MBCs in blood and tonsils of both vaccinated and recovered individuals, whereas nucleocapsid^+^ MBCs were only enriched in blood and tonsils of subjects that had contact with the virus (Fig. 3a,b). Spike^+^ tonsillar MBCs showed slightly lower percentages of IgG and IgM positivity, but IgA^+^ cells were more frequent in tonsils than in circulation (Fig. 3c). Analyzing SARS-CoV-2-specific Bcl-6^+^Ki-67^+^ GC B cells, we found a trend toward elevated levels of spike^+^ and nucleocapsid^+^ GC cells in subjects recovered from COVID-19 compared to individuals only vaccinated (Supp. Fig. 4c).

**Fig. 3.**
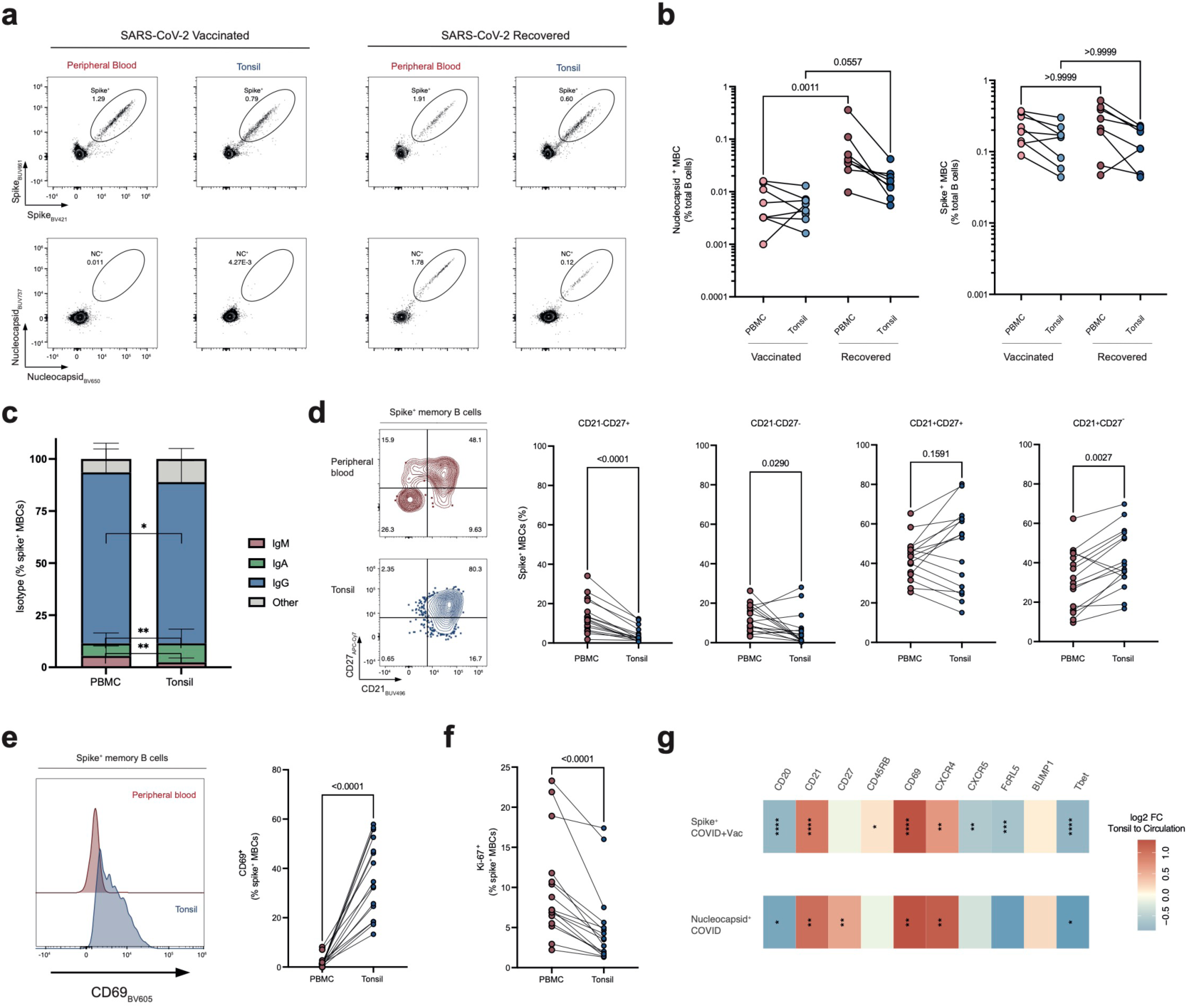
Phenotypes of circulating and tonsillar SARS-CoV-2-specific MBCs upon infection and vaccination. **a**, Representative plots MBCs, showing spike^+^ (top) and nucleocapsid^+^ (bottom) cells in a vaccinated (CoV-Tissue-01; left) and COVID-19-recovered (CoV-Tissue-02; right) individual in paired tonsil and peripheral blood samples. **b**, Nucleocapsid^+^ (left) and spike^+^ (right) MBC frequencies in peripheral blood and paired tonsil samples. Lines connect paired samples. Groups are separated by SARS-CoV-2 vaccination (n=8) and recovered status (n=8). Frequencies in same compartment between different groups were compared. **c**, Stacked histograms showing isotype distribution in spike^+^ MBCs in peripheral blood and paired tonsils, mean + SD. Samples from vaccinated and COVID-19-recovered individuals were combined for analysis (n=16). **d**, Contour plots of spike^+^ MBCs of peripheral blood (red, top left) and tonsil (blue, bottom left) of patient CoV-Tissue-02. Frequencies of indicated subsets within spike^+^ MBCs in peripheral blood and paired tonsils (right). Lines indicate paired samples. Samples from vaccinated and COVID-19-recovered individuals were combined for analysis (n=16). **e**, Representative histograms of CD69 (CoV-Tissue-02, left) and percentages of CD69^+^ spike^+^ MBCs in peripheral blood and tonsils (right). **f**, Percentages of Ki-67^+^ spike^+^ MBCs in peripheral blood and tonsils. **g**, Heatmap of log2-fold change of indicated markers in peripheral blood and tonsils, with red indicating higher expression in tonsils and blue in peripheral blood, in spike^+^ MBCs (top) of vaccinated and COVID-19-recovered individuals (n=16) and in nucleocapsid^+^ MBCs (bottom) of COVID-19-recovered individuals (n=8). Unpaired samples were compared with Mann-Whitney test, paired tests with Wilcoxon matched-pairs signed rank test. In case of more than two groups, unpaired testing was performed using Kruskal-Wallis test with Dunn’s multiple comparison correction. In **b** and **d**, all p-values are shown, in other plots p-values are shown if significant (p<0.05). *p<0.05, **p<0.01, ***p<0.001, **** p<0.0001

Assessing spike^+^ MBC subsets, CD21^−^CD27^+^ activated and CD21^−^CD27^−^ atypical MBCs were found at higher frequencies in blood, whereas CD21^+^ resting MBCs were more abundant in tonsils (Fig. 3d). Compared to their circulating counterparts, tonsillar spike^+^ and nucleocapsid^+^ MBCs expressed, on average, higher CD69, lower Ki-67, lower T-bet, and different chemokine receptor levels (Fig. 3e–g), suggestive of a resting and tissue-resident memory phenotype. Altogether, these data suggested that SARS-CoV-2 infection and mRNA vaccination led to the induction of long-lived and resting antigen-specific MBCs, which homed to peripheral secondary lymphoid organs where they acquired characteristics of tissue residency.

### Changes in MBC subsets following vaccination

With the availability of SARS-CoV-2 mRNA vaccines, 35 individuals of our COVID-19 cohort got vaccinated between the six- and 12-month and three subjects before the six-month sampling time point (Fig. 1a, Supp. Table 1). This setting allowed us to investigate the MBC response to controlled antigen reexposure. Vaccination resulted in approximately five-fold increase in circulating MBC frequencies (Fig. 4a,b). As sampling occurred at different time points after vaccinations, time-resolved analysis of spike^+^ MBCs revealed an early peak after vaccination followed by a slow decrease in frequencies (Fig. 4c). In our scRNA-seq dataset, we observed an increased clonality in paired samples after vaccination (Fig. 4d). Moreover, we found counts of SHM remained high in spike^+^ MBCs even after vaccination compared to six-month samples (Fig. 4e).

**Fig. 4.**
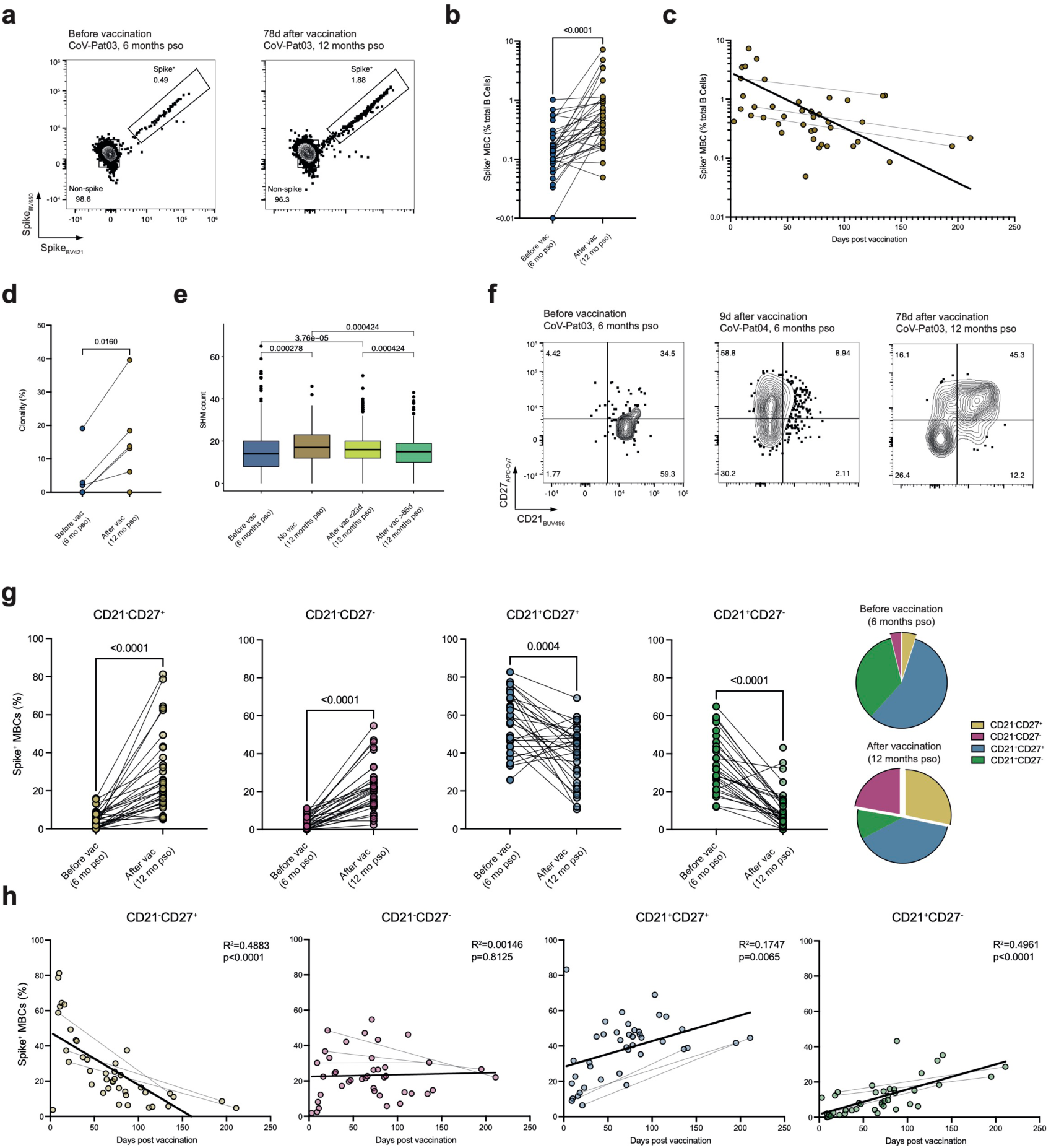
Changes in antigen-specific MBC subsets following vaccination. **a**, Representative flow cytometry plots of spike^+^ MBCs before (left; six months after acute infection) and 78 d after (right; 12 months after infection) vaccination (vac) in the same patient (CoV-Pat03). **b**, Paired comparison of spike^+^ MBC frequencies (n=34) before (six months after infection) and after vaccination. **c**, Spike^+^ MBC frequencies (n=41) plotted as time after last vaccination, with lines connecting paired samples. Semilog line fitted to data (R^2^=0.2695). **d**, Clonality analysis of spike^+^ MBCs before (six months after infection) and after vaccination. Each dot represents one individual (n=6). **e**, SHM counts of spike^+^ MBCs before (n=9; six months), without (n=3; 12 months), as well as early (n=3; less than 23 d) and late (n=3; more than 85 d) after vaccination. **f**, Representative flow cytometry plots of CD21 and CD27 on spike^+^ MBCs before and early and late after vaccination. **g**, Frequencies of spike^+^ MBC subsets (n=29) and pie chart distribution (far right) of indicated MBC subsets at indicated time points. **h**, Percentages of spike^+^ MBC subsets plotted as time after last vaccination. Lines combine paired samples. Linear regressions are fitted to data. Paired samples were compared with a Wilcoxon matched-pairs signed rank test (**b**, **g**) or paired t-test (**d**). In **e**, two-sided Wilcoxon test was used. Holm-Bonferroni method was used for p-value adjustment of multiple comparisons.

Whereas spike^+^ MBCs were predominantly of a resting CD21^+^ memory phenotype at six months, SARS-CoV-2 mRNA vaccination very strongly induced the appearance of spike^+^ CD21^−^CD27^+^ activated and CD21^−^CD27^−^ atypical MBCs in blood (Fig. 4f,g). Spike^+^ CD21^−^CD27^+^ activated MBCs sharply peaked after vaccination followed by a rapid decline thereafter. Conversely, frequencies of spike^+^ CD21^−^CD27^−^ atypical MBCs, which accounted for about 20% of spike^+^ MBCs, remained stable after vaccination (Fig. 4h).

### Transcriptional makeup of SARS-CoV-2-specific MBC subsets

To gain insight into pathways guiding development of different MBC subsets we focused on our scRNA-seq dataset of sorted spike^+^ and spike^−^ memory B cells (Supp. Fig. 5a). Based on weighted-nearest neighbor (WNN) clustering of all sequenced MBCs, we identified 10 clusters and subsequently merged these into five subsets based on surface markers, determined by oligonucleotide-tagged antibodies, and isotype expression (Supp. Fig. 5b). We annotated these five subsets as CD21^−^CD27^+^CD71^+^ activated, CD21^−^CD27^−^FcRL5^+^ atypical, CD21^+^CD27^−^ resting, CD21^+^CD27^+^ resting, and unswitched MBCs (Fig. 5a, Supp. Fig. 5b). We subsequently focused on the spike^+^ MBCs. Six and 12 months after acute SARS-CoV-2 infection the predominant subset in individuals not receiving vaccination consisted of CD21^+^ resting MBCs, whereas activated and atypical MBCs made up the main subsets in subjects at 12 months after infection that had been vaccinated (Fig. 5a,b, Supp. Fig. 5d). As expected, unswitched MBCs were almost entirely IgM^+^, whereas the other MBC subsets expressed mainly IgG subclasses, with atypical MBCs containing the lowest fraction of IgM^+^ cells (Supp. Fig. 5e). These findings were consistent with our flow cytometry analysis of Ig subclasses in spike^+^ MBCs (Supp. Fig. 5f).

**Fig. 5.**
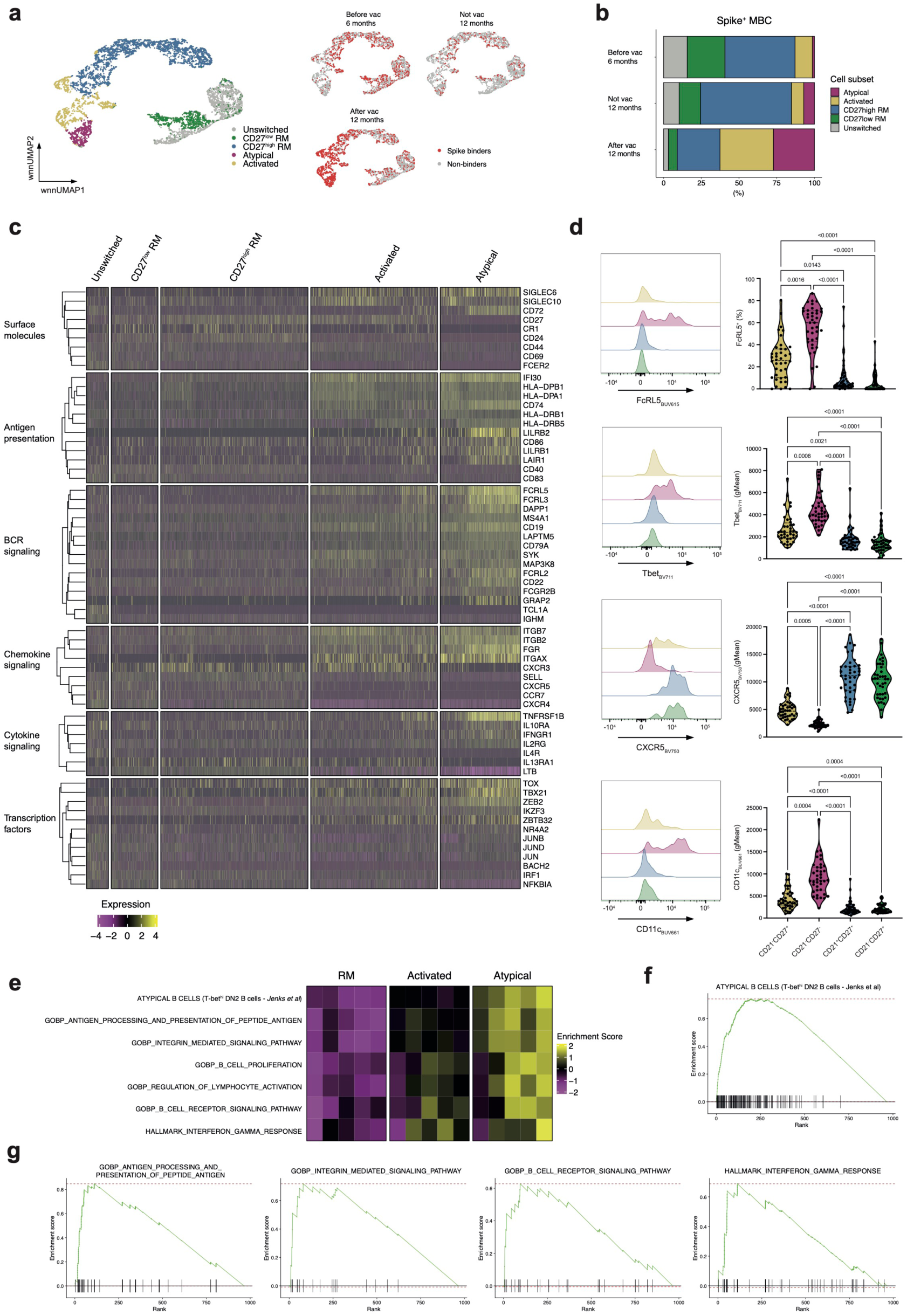
Transcriptional makeup of SARS-CoV-2-specific MBC subsets. **a**, Weighted-nearest neighbor UMAP (wnnUMAP) analysis of indicated MBC subsets (n=9; left) and of MBCs binding or not binding to spike (right) at six months and 12 months after infection without vaccination, and at 12 months after infection and with vaccination. **b**, Distribution of spike^+^ MBC subsets at indicated time points. **c**, Heatmap of selected, significantly differentially expressed genes in MBC subsets. Functional groups of genes, ordered by hierarchical clustering. **d**, Representative histograms (left) and violin plots of indicated markers on spike^+^ MBC subsets (right; n=41) after vaccination. Markers were compared if a subset had more than 3 cells. **e**, Heatmap of enrichment scores of selected gene sets, comparing CD21^+^ resting memory (RM), activated, and atypical spike^+^ MBCs in a pseudo bulk analysis (n=5 individuals, patients with more than 20 cells in each MBC subset). **f–g**, Gene set enrichment analysis of atypical versus RM spike^+^ MBCs for selected gene sets. Red dashed lines indicate minimal and maximal cumulative enrichment values. Samples in **d** were compared using Kruskal-Wallis test with Dunn’s multiple comparison correction. Adjusted p-values are shown if significant (p<0.05).

Analysis of significant differentially-expressed genes (DEGs) revealed marked differences in the spike^+^ MBC subsets in terms of transcription factors, signaling, surface molecules, and antigen presentation (Fig. 5c). Atypical MBCs were the most distinctive subset, expressing the highest levels of *TBX21* (encoding T-bet), the T-bet-driven genes *ZEB2* and *ITGAX*, and *TOX*. Moreover, atypical MBCs were enriched in gene transcripts involved in interferon-γ and BCR signaling and showed high expression of the integrins *ITGAX*, *ITGB2*, and *ITGB7*. Notably, expression of inhibitory receptors, including *FCRL2*, *FCRL3*, *FCRL5*, *SIGLEC6*, *SIGLEC10*, *LAIR1*, *LILRB1*, and *LILRB2*, was particularly high in atypical MBCs. Furthermore, atypical MBCs showed high expression of proteins involved in antigen presentation and processing, such as *HLA-DPA1*, *HLA-DPB1*, *HLA-DRB1*, *HLA-DRB5*, *CD74*, and *CD86*. Several of these differences were also confirmed on protein level (Fig. 5d).

To further investigate differentially regulated processes in the subsets we performed gene set variation and enrichment analysis, respectively (Fig. 5e-g). Using a previously described atypical B cell signature^44^, we found a strong enrichment of this signature in our respective SARS-CoV-2-specific MBC subset. In line with the DEGs, gene sets involved in antigen presentation and integrin-mediated signaling were prominently enriched in atypical MBCs compared to resting and activated MBCs (Fig. 5e). Moreover, also B cell activation, BCR and interferon-γ signaling was highly upregulated in atypical MBCs (Fig. 5e–g). All these transcriptional hallmarks of atypical MBCs were very differently expressed in the other MBC subsets (Fig. 5c–e).

### Clonal relationships between antigen-specific MBC subsets

Our setup allowed us to longitudinally track spike^+^ MBC clones to investigate the relationship of the different MBC subsets and the factors guiding their fate decisions. Indeed, we observed some clonal overlap between spike^+^ resting, activated, and atypical MBCs (Supp. Fig. 6a). Subsequently, we focused on the longitudinal aspect and identified 30 persistent clones between six and 12 months after SARS-CoV-2 infection in individuals vaccinated during that period (Fig. 6a, Supp. Fig. 6b). In individuals at six months after acute infection and before vaccination, about 80% of persistent MBC clones were of a CD21^+^ resting phenotype (Fig. 6b). Conversely, upon vaccination, about 33% of MBC clones showed an activated and another 30% an atypical phenotype, both in persistent and newly detected clones (Fig. 6b).

**Fig. 6.**
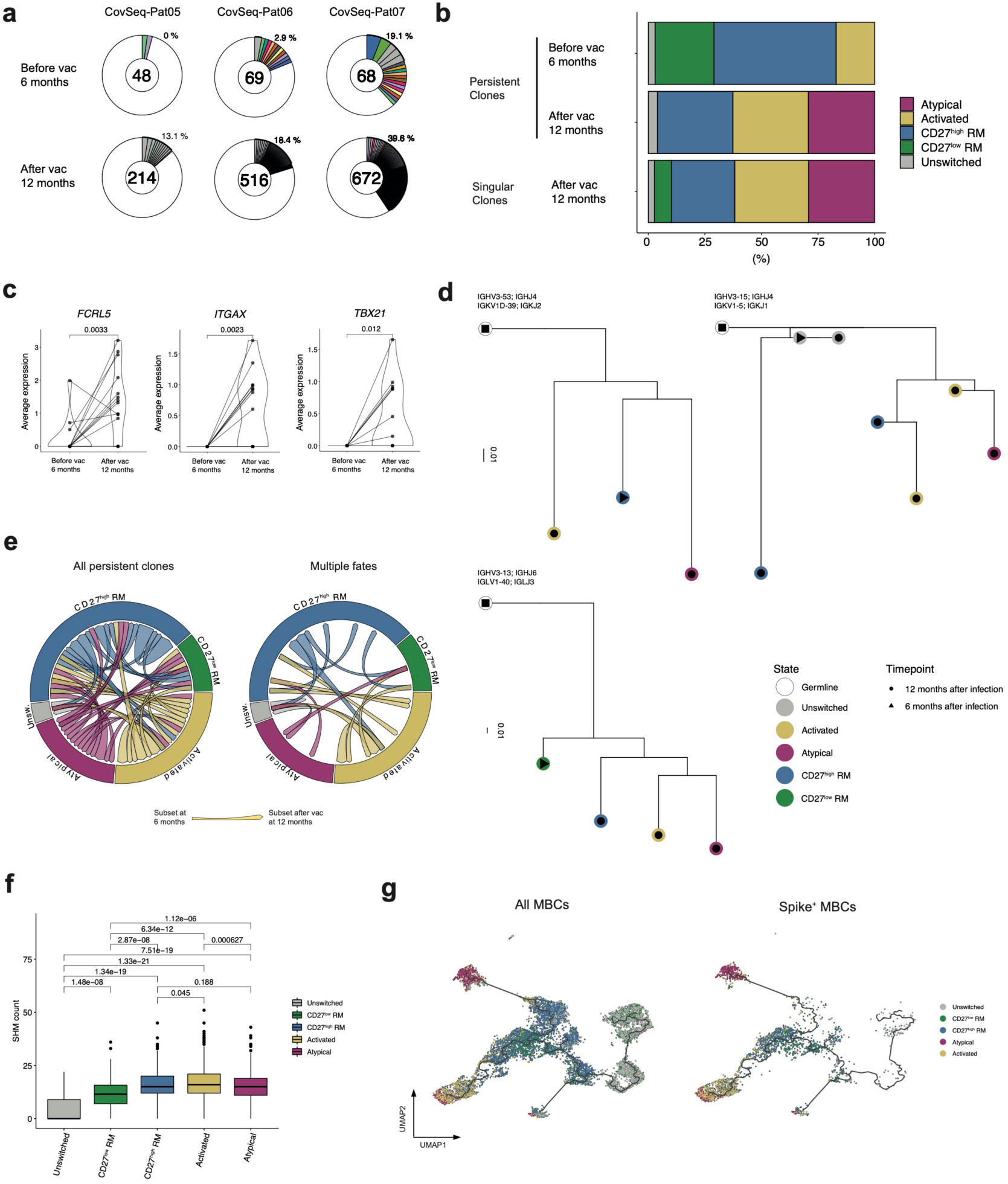
Temporal analysis of B cell receptors in SARS-CoV-2-specific MBC subsets. **a**, Donut plots of B cell receptor (BCR) sequences of spike^+^ MBCs in three representative patients before and after vaccination. Numbers in donuts indicate spike^+^ MBCs. Gray slices indicate singular clones found at one time point only, whereas persistent clones found at both six and 12 months after infection are labeled by the same color. White areas represent BCR sequences found in single cells only. Slice sizes correspond to clone sizes. Percentages indicate frequencies of clonally expanded cells. **b**, Distribution of spike^+^ MBC subsets in persistent and singular clones at indicated time points. **c**, Average expression of indicated genes – before and after vaccination – in persistent clones of spike^+^ MBCs that contained at least one atypical MBC. **d**, Exemplary dendrograms (IgPhyML B cell trees) of different MBC clones at six (dots) and 12 months (triangles) after infection. Colors indicate MBC subsets. Germline sequences, inferred during the Immcantation pipeline, are shown in white. Branch lengths represent mutation numbers per site between each node. VH and VL genes of clones are indicated on top of each dendrogram. **e**, Circos plot of persistent spike^+^ MBC clones, with arrows connecting cells of six-with 12-month time points and coloring according to MBC phenotype at 12 months. All clones (left) versus clones adopting multiple MBC fates (right) are shown. **f**, SHM counts of indicated spike^+^ MBC subsets. **g**, UMAP representation of Monocle 3 analysis of all (left) and spike^+^ MBCs (right). Colors indicate MBC subsets. Black lines indicate trajectory. Samples were compared using paired t test (**c**) or two-sided Wilcoxon test (**f**). Holm– Bonferroni method was used for p value adjustment of multiple comparisons.

We could demonstrate by three different measures that cells of individual MBC clones were able adopt different MBC fates following vaccination at six to 12 months after infection. Firstly, we saw in single MBC clones upregulation of genes associated with the atypical B cell lineage, including *TBX21*, *ITGAX* and *FCRL5* (Fig. 6c). Secondly, by reconstructing clonal lineage trees, we found that cells of individual MBC clones acquired different MBC fates (Fig. 6d). And thirdly, we obtained similar results when visualizing persistent MBC clones in a circos plot, showing that cells of a given MBC clone could adopt different MBC phenotypes (Fig. 6e).

Assessment of SHMs in the different spike^+^ MBC subsets revealed that only unswitched and CD27^−^ resting MBCs had low counts, whereas CD27^+^ resting, activated, and atypical MBCs contained comparably high counts (Fig. 6f). To further investigate the connection of MBC subsets, we performed a pseudotime-based trajectory analysis using Monocle 3 with our scRNA-seq dataset (Supp. Fig. 6c–d). Visualizing the cell subsets from our previous analysis on the Monocle UMAP space, we identified two branches, which strongly separated activated and atypical MBCs and branched out from the resting MBCs (Fig. 6g, Supp. Fig. 6e). Altogether, these findings show that upon immunization in SARS-CoV-2 recovered individuals, antigen-specific MBCs acquired distinct clonal differentiation fates along different trajectories.

## DISCUSSION

In this study, we demonstrated that individual MBC clones harbored the capacity to adopt multiple and functionally different fates in COVID-19 patients during immunological memory and SARS-CoV-2-specific immunization. Thus, single MBC clones were able to differentiate in vivo into CD21^−^CD27^+^ activated, CD21^−^CD27^−^ atypical, or CD21^+^CD27^+/–^ resting MBCs upon vaccination. Whereas activated MBCs peaked and declined rapidly after vaccination, atypical MBCs remained stable and resting MBC subsets even increased in percentages. Moreover, we found an enrichment of SARS-CoV-2-specific resting MBCs in peripheral lymphoid organs carrying features of tissue-resident cells. Overall, these data provide evidence that single MBC clones can give rise to diverse MBC trajectories with phenotypically and functionally different characteristics during recall, including atypical MBCs.

Atypical MBCs have been previously observed in chronic infections with *Plasmodium falciparum*, human immunodeficiency virus (HIV), and hepatitis C virus, in immunodeficiencies, as well as in autoimmune diseases^21, 22, 25, 26, 45^. Moreover, these cells share certain features of so-called age-associated B cells found in mice, which are also characterized by high expression of CD11c and T-bet and implicated in autoimmunity^46, 47^. Recently, influenza-specific atypical MBCs have been described transiently during de novo, but not recall, influenza vaccine responses^16^, as well as during acute SARS-CoV-2 infection and vaccination^27, 31–35^, the latter of which is in line with our findings. Intriguingly, we find that SARS-CoV-2-specific atypical MBCs are transcriptionally very similar to their counterparts in autoimmune disease. These results further confirm that atypical MBCs are part of the normal immune response against different pathogens^48^.

A defining feature of atypical MBCs appears to be increased BCR and interferon-γ signaling, which likely induce and govern their T-bet-controlled program. In line with this suggestion, a recent study elegantly demonstrated that efficient T-bet expression in human B cells required strong BCR and interferon-γ receptor stimulation along with signals from pathogen-associated molecular patterns or from Th cells^38^. Confirming and extending these data, we also found in SARS-CoV-2-specific atypical MBCs signs of increased BCR and interferon-γ signaling, the latter of which fitted well with the increased levels of T-bet and the T-bet target genes *ZEB2* and *ITGAX* (encoding CD11c). Conversely, atypical B cells were completely absent in a patient with an inborn T-bet deficiency^37^, demonstrating the crucial role of T-bet also in human atypical B cells.

Several models have been proposed to explain the development of MBC heterogeneity^49, 50^. Thus, MBC subsets could comprise entirely separate lineages with different BCR repertoires or single B cell clones could give rise to phenotypically and functionally different MBC subsets, with stably imprinted phenotype and function versus full or hierarchical plasticity. Our longitudinal data are in line with the latter model in showing that different MBC subsets were clonally related. Studies in humans with systemic lupus erythematosus or following HIV infection suggest that atypical MBCs differentiated via an extrafollicular pathway, thus avoiding GCs^23, 24^. In our analysis, the SHM counts in different antigen-specific MBC subsets revealed that resting, activated, and atypical MBCs contained comparable levels SHMs. This could either reflect a GC origin of the subsets including atypical MBCs or that atypical MBCs originate from a GC-derived progenitor MBC upon antigen rechallenge. The latter hypothesis fits well with our clonal analysis. It remains to be investigated whether the atypical MBCs we observed following vaccination in the memory phase can again become resting MBCs or whether their functional phenotype remains fated. The expression of *ZEB2* in atypical MBCs could suggest the latter, as ZEB2 together with T-bet commits CD8^+^ effector T cells to a terminal differentiation state and has been proposed to act similarly in B cells^23, 51^.

Whether atypical MBCs contribute to protective immunity in acute and chronic infection in humans remains a field of controversy^52^. T-bet-expressing B cells, including atypical MBCs, played a protective role in mouse models of acute and chronic viral infections^49, 53^. Moreover, intrinsic T-bet expression in B cells was essential for forming long-lived antibody-secreting B cells in a mouse model of influenza infection^54^. However, in the above-mentioned T-bet-deficient patient, antibody responses to several previously-applied vaccines, such as tetanus toxoid, diphtheria, *Haemophilus influenzae* type b, and pneumococcal antigen, were all normal^37^. Atypical MBCs in humans were thought to have a reduced capacity to develop into antibody-secreting cells due to their expression of inhibitory receptors, albeit recent studies showing they can secrete antibodies when receiving T cell help and act as antigen-presenting cells^29, 55^. According to our results, spike^+^ atypical MBCs carried signs of increased antigen processing and presentation, however, a recent study in atypical B cells isolated from patients with systemic autoimmune diseases failed to show an enhanced ability of these cells to stimulate Th cells in vitro compared to other B cell subsets^56^.

Our data showed that SARS-CoV-2 infection induced long-lived, stable antigen-specific MBCs in the circulation^57, 58^. These cells acquired a CD21^+^ resting memory phenotype in the memory phase and continued to mature up to one year after infection, as evidenced by their elevated proliferation rate, increasing SHM counts, and improved breadth of SARS-CoV-2 antigen recognition. This is in line with previous publications showing that SARS-CoV-2 infection led to lasting MBC maturation via an ongoing GC reaction, potentially due to persistent antigen^32, 59, 60^. COVID-19 severity did not appear to significantly affect frequencies of SARS-CoV-2-specific MBCs, except for a tendency to more spike^+^ MBCs in severe COVID-19, which is consistent with previous findings of increased spike-specific Ig in these patients^61^. These observations in circulating MBCs were paralleled by the appearance of resting MBCs we found in tonsils where they showed high expression of CD69, low proliferation rates, and low levels of T-bet. The phenotype, together with the chemokine receptor expression of the different subsets in the circulation is suggestive that these cells arise from the CD21^+^ resting MBC subsets. Previous work found an enrichment of hemagglutinin-specific atypical MBCs in the human spleen^49^. Considering the chemokine receptor profile of atypical MBCs it is intriguing to speculate that the cells could migrate to tissue niches^62^. CD69 expression is a hallmark of tissue residency in T cells^4, 63^ and has also been proposed to characterize resident memory B cell populations in lymphoid and non-lymphoid human tissues^64, 65^. A previous publication found SARS-CoV-2-specific CD69^+^ MBCs in lungs and in lung- and gut-draining lymph nodes of COVID-19 recovered individuals^66^. Unfortunately, we were unable to receive other tissue samples other than tonsils to extend our findings.

Other potential shortcomings of our study include the limitation that our clonal analysis was restricted to the vaccination setting, as cell numbers during acute infection were too low for our sequencing approach. Moreover, although our multimer staining approach has been previously used^57, 58, 67^ and we further confirmed the validity by identifying several previously described variable chains to be enriched in RBD^+^ MBCs^40–42^, this approach might miss low-affinity antigen binders^68^. Based on our data, we favor a linear-plastic model where stimulation and GC maturation of antigen-specific B cells results in MBCs that gradually adopt a CD21^+^ Ki-67^low^ bona fide resting state between 6-12 months after acute infection. These resting MBCs may circulate in the blood, thus providing a mobile unit of primed B cells able to rapidly respond to antigen rechallenge upon which they can acquire different MBC fates. Or they might home to secondary lymphoid and peripheral organs where they form a CD69^+^ tissue-resident defense line, ready to deploy at potential entry sites of pathogens. Although they are currently unknown for MBCs, identification of the signals instructing resting MBCs to migrate to peripheral sites might guide immunization and booster strategies aimed at tissues and certain MBC subsets. On the latter, our work sheds further light on atypical MBCs, which appear to make up a sizeable portion of MBCs following an acute viral infection and vaccination in humans.

## METHODS

### Flow cytometry cohort and scRNA-seq subcohort

Following written informed consent COVID-19 patients were recruited at four hospitals in the Canton of Zurich, Switzerland. The study was approved by the Cantonal Ethical Committee of Zurich (BASEC #2016-01440). Patients had to have a reverse-transcriptase polymerase chain reaction (RT-PCR) confirmed SARS-CoV-2 infection and be symptomatic to be included in the study. Subsequently, patients visited again at six and at 12 months after infection and donated blood and serum samples at the respective time points, which was processed and biobanked. The full cohort and biobanking process has been previously described^69, 70^. We included a total of 65 patients, 42 with mild COVID-19 and 23 with severe COVID-19, from the full cohort (and 1 healthy control which was infected subsequently to establish the staining specificity) based on a power calculation from pre-experiments. Patients were selected according to the sample availability and had to have at least paired samples from 2 time points. The full flow cytometry cohort and scRNA-seq subcohort characteristics are shown in Supp. Table 1 and 2 respectively. The patients were included in the study during their acute disease between April 2020 and September 2020 and for the 12 months follow-up between April 2021 and September 2021.

### Tonsil cohort

Paired tonsil and peripheral blood samples, as well as serum samples, were collected from patients undergoing a tonsillectomy at the University Hospital Zurich between November 2021 and April 2022. All patients signed a written informed consent (BASEC #2016-01440) before sample collection. Patients underwent their tonsillectomy for recurrent and chronic tonsillitis or obstructive sleep apnea. Clinical data regarding SARS-CoV-2 infection and vaccination was evaluated from history and derived from electronic medical records. From all the patients SARS-CoV-2 spike and nucleocapsid specific antibodies were measured (see below), patients were assigned as “SARS-CoV-2 recovered” if they had a confirmed SARS-CoV-2 infection and/or SARS-CoV-2 nucleocapsid antibodies. The cohort size was based on sample availability. The full cohort characteristics are shown in Supp. Table 3.

Peripheral blood and serum were processed and biobanked as described. Tonsils were processed according to established protocols^66, 71^. Briefly, they were mechanically cut into smaller pieces, grinded through a 70 μm cell strainer, washed in phosphate buffered saline, before a density gradient centrifugation was performed. Subsequently, the mononuclear cells were washed, counted, frozen in fetal bovine serum with 10% dimethyl sulfoxide (DMSO) and stored in liquid nitrogen until use.

### Spectral flow cytometry

To stain antigen-specific B cells, we probe multimers were created, similarly, to previously described protocols^58, 67^. Commercially available biotinylated SARS-CoV-2 spike, RBD, nucleocapsid (MiltenyiBiotec) and H1N1 (A/California/07/2009, SinoBiological) were incubated separately with fluorescently labelled streptavidin (SAV) at 4:1 molar ratio for SARS-CoV-2 proteins and 6:1 for influenza antigen. SAV was added stepwise every 15 min at 4°C for 1hr. Subsequently, the staining mix was created by mixing the probes in 1:1 Brilliant Buffer (BD Bioscience) and FACS buffer (PBS with 2% FBS and 2mM EDTA) with 5μM of free D-biotin. For the tonsil cohort staining spike (separate multimers with SAV-BUV661 and BV421), RBD (SAV-KIRAVIA520), nucleocapsid (separate multimers with SAV-BUV737 and BV650), and hemagglutinin (SAV-BV785) and for the full cohort staining spike (separate multimers with SAV-BV421 and SAV-BV650), RBD (SAV-PE/Cy7) and a decoy probe (SAV-BV785) were combined, termed “antigen-specific stain mix”.

Frozen mononuclear cells (∼5×10^6^ cells) were thawed and plated in 96 U-bottom well plates. They were then stained with ZombieUV Live-Dead staining (1:400, Biolegend) and TruStain FcX (1:200, Biolegend) in PBS for 30 min, washed with FACS buffer and subsequently stained with 50 μl of staining mix with the antigen-specific stain mix (200 ng spike, 50 ng RBD, 100 ng nucleocapsid, 20 ng SAV-Decoy per color per 50 μl) for 1 hr at 4°C. After washing, the cells were then stained for 30 min with the surface staining mix. The cells were then fixed and permeabilized with 200 μl transcription factor staining buffer (eBioscience) at room temperature for 1 hr. Lastly, cells were stained intracellularly with the intracellular staining mix in PermWash for 30 min at room temperature, before washing and resuspending in FACS buffer for acquisition (for full staining see Supp. Tables 4 and 6). The staining mixes were centrifuged at 14000 g for 2 min before staining. Subsequently, the samples were acquired on a Cytek Aurora spectral flow cytometer using the SpectroFlo software. Quality control for the cytometer was performed daily. The samples were analysed in several batches, paired samples were always recorded in the same batch. Furthermore, in every experiment the same positive control from a SARS-CoV-2 vaccinated healthy control was included to ensure consistent results.

### SARS-CoV-2 antibody measurement

For the full patient cohort, the anti-SARS-CoV-2 antibodies were measured by a commercially available enzyme-linked immunosorbent assay (ELISA) specific for the S1 protein of SARS-CoV-2 (Euroimmun SARS-CoV-2 IgG and IgA) as previously described^61^. In the tonsil cohort, the IgG, IgA and IgM response against SARS-CoV-2 RBD, S1, S2 and N was measured with a bead-based multiplexed immunoassay available at the Department of Medical Virology from the University of Zurich termed AntiBody CORonavirus Assay (ABCORA) which has previously been described^72^.

### Cell Sorting for scRNA-seq

To sort SARS-CoV-2 specific and non-specific memory B cells samples were processed similarly as for the spectral flow cytometry staining. Briefly, commercially available biotinylated SARS-CoV-2 spike (spike_WT_), RBD, spike variants beta and delta (MiltenyiBiotec) were multimerized as described with fluorescently labelled and/or barcoded streptavidin (TotalSeqC, Biolegend) (4:1 molar ratio). The total antigen-staining mix contained: spike_WT_ separate multimers with SAV-BV421 and barcoded SAV-PE; RBD SAV-barcoded; spike beta variant barcoded SAV-PE, spike delta variant barcoded SAV-PE and separate decoy probe SAV-BV785 and SAV-barcoded. Frozen PBMCs were thawed, stained in a 96-U bottom well plate with fixable viability dye eFluor^TM^780 (eBioscience) and TruStainFcX for 20 min at 4°C, washed and then stained for 1hr with the antigen-specific stain mix. After washing, cells were stained for 30 min at 4°C with a surface staining mix which contained fluorescently labelled antibodies and a panel of barcoded antibodies (CD21, CD27, CD71, CXCR5, FcRL5; in the sample set where naïve B cells were sorted IgD), also each sample was stained with a hashtag antibody for sample multiplexing (for full panel see Supp. Table 5). After 3 washing steps, the cells were resuspended and sorted on a FACS Aria III 4L sorter using a 70-μm nozzle. All specific cells per sample were sorted together with 1500-2000 non-specific memory B cells. In one sample set we additionally sorted naïve B cells (500 per sample). Cells were sorted into the same tube.

### scRNA-seq sequencing and library preparation

FACS-sorted B cells were analyzed by single cell RNA sequencing (scRNA-seq) utilizing the commercial 5′ Single Cell GEX and VDJ v1.1 platform (10x Genomics). After sorting, cell suspensions were pelleted at 400 g for 10 min at 4°C, resuspended and loaded into the Chromium Chip following the manufacturer’s instructions. 14 cycles (in one case 17) of initial cDNA amplification were used for all sample batches and single-cell sequencing libraries for whole-transcriptome analysis (GEX), BCR profiling (VDJ), and TotalSeq (Biolegend) barcode detection (ADT) were generated. Final libraries were quantified using a Qubit Fluorometer, pooled in a ratio of 5:1:1 or 10:1:1 (GEX:VDJ:ADT) and sequenced on a NovaSeq 6000 system.

### Flow Cytometry Analysis

Flow cytometry data were analysed with FlowJo (version 10.8.0), full gating strategies are shown in Figures S1, S4. Subsets and markers of antigen-specific B cells were evaluated only in patients with >9 specific cells per sample and of antigen-specific subsets only if the subset had >3 specific cells. Dimensionality reduction and clustering analysis of flow cytometry data was performed using OMIQ (www.omiq.ai). Markers were scaled with an arcsinh-transformation (cofactor 6000), the samples were subsetted to maximally 25 spike^+^ MBCs per sample. For UMAP representations and PhenoGraph clustering, k was set to 20^73^ and B cell markers of interest were used, including CD11c, CD19, CD20, CD21, CD24, CD27, CD38, CD71, CD80, CXCR5, BAFF-R, FcRL5, IgA, IgD, IgG, IgM, Blimp1, IRF8, Ki67, and Tbet.

### Single-cell transcriptome analysis

Preprocessing of raw scRNA-seq data was done as described before^69^. Briefly, FASTQ files were aligned to the human GRCh38 genome using Cell Ranger’s ‘cellranger multi’ pipeline (10x Genomics) (v6.1.2) with default settings, which allows to process together the paired GEX, ADT and VDJ libraries for each sample batch. Downstream analysis was conducted in R version 4.1.0 mainly with the package Seurat (v4.1.1)^74^. Cells with fewer than 200 or more than 2,500 detected genes and cells with more than 10% detected mitochondrial genes were excluded from the analysis. Gene expression levels were log normalized using Seurat’s NormalizeData() function with default settings. Sample assignment of cells was done using TotalSeq based cell hashing and Seurat’s HTODemux() function. When comparing dataset quality, we noticed a markedly lower median gene detection and UMI count per cell in one of our datasets. We associated this with an incident during sample preparation in one of our experiments and decided to exclude most cells of this dataset from the analysis.

As an internal reference for SHM counts in naïve B cells, we co-sorted naïve B cells in one of our experiments. Before integration of this dataset with others and doing further downstream analysis, we excluded these cells from the dataset. For this, cells were clustered alone and naïve B cell clusters were identified based on their surface protein expression levels of CD27, CD21 and IgD as well as on their RNA levels of naïve B cell markers *TCLA1*, *IL4R*, *BACH2*, *IGHD* and *BTG1*. Independent datasets were then integrated using Seurat’s anchoring-based integration method. Gene expression data and TotalSeq surface proteome data were first integrated separately. Then, Seurat’s weighted nearest neighbor analysis was used to take advantage of our multimodal approach during clustering and visualization^74^. Clustering was performed using the Louvain algorithm and a resolution of 0.4. For UMAP generation, the embedding parameters were manually set to a=1.4 and b=0.75. Differential gene expression analyses were done using assay ‘RNA’ of the integrated dataset. FindAllMarkers and FindMarkers functions were executed with logfc.thresholds set to 0.25 (0.1 when comparing RM cells at six months versus 12 months) and a min.pct cutoff at 0.1. Heatmaps were generated using the ComplexHeatmap package (v2.13.1)_75._

Gene set enrichment analysis (gsea) was done as described before^69^. Briefly, lists of differentially expressed genes were first pre-ranked in decreasing order by the negative logarithm of their P value, multiplied for the sign of their average log-fold change (in R, ‘-log(P_val)*sign(avg_log2FC)’). Gsea was then performed on this pre-ranked list using the R package fgsea (v.1.2). Gene sets were obtained from the Molecular Signatures Database (v7.5.1, collections H and C5) and loaded in R by the package msigdbr (v.7.5.1). To make the results reproducible, the seed value was set (‘set.seed(42)’ in R) before execution. fgsea uses a P value estimation based on an adaptive multi-level split Monte-Carlo scheme. A multiple hypothesis correction procedure was applied to get adjusted P values. Finally, results were filtered for gene sets that were significantly enriched with adjusted P < 0.05.

Gene set variation analysis with the package gsva (v1.42.0) was used to estimate gene set enrichments for more than two groups^76^. Transcriptomes of individual cells were used as inputs for the gsva() function with default parameters. Then, gene set enrichments for individual cells were summarized to patient pseudo bulks by calculating the mean enrichment values of cells belonging to the same patient. Pseudo bulking was only done for patients with n>20 cells in each cell subset. The resulting scores were used to compute fold changes and significance levels for enrichment score comparisons between cell subsets in limma (v3.50.3)^77^.

Single cell trajectories were created with Monocle3 (version 1.2.9)^78^. First, raw counts obtained from the cellranger gene expression matrix were used to create cell data sets, which were then pre-processed using the Monocle 3 pipeline. Different batches were aligned using Batchelor (v.1.10.0)^79^. The num_dim parameter of Monocle’s preprocess_cds() function was set to 20. Functions reduce_dimension(), order_cells() and graph_test() were executed with default parameters.

### BCR analysis

B cell clonality analysis was performed mainly with the changeo-10x pipeline from the Immcantation suite^80^ using the singularity image provided by Immcantation developers. filtered_contig_annotations.csv files obtained from the cellranger multi pipeline were used as input for the changeo-10x pipeline. Unique combinations of bases were appended to the cell barcodes per batch before combining the data from different batches of sequencing to prevent cell barcode collisions. The clonality distance threshold was set to 0.20. Visualization of the clonal trees was done using dowser^81^. BCR variable gene segment usage was additionally quantified using the R package scRepertoire (v.1.3.5)^82^.

### Mapping of B Cell Receptor Sequences to Antigen Specificity

We used an adaptation of LIBRAseq^83^ to identify antigen specific cells in our sequencing data. First, the raw counts from the baiting negative control were subtracted from the counts of all other antigen-baiting constructs in every cell. Then, cutoffs for background binding levels were manually determined for every construct individually by inspection of the bimodal distribution of count frequencies across all cells. All binding counts falling below thresholds were set to zero and hence classified as “non-binding”. Next, Seurat’s centered log ratio transformation was applied across features, followed by a scaling of the obtained values. This resulted in final LIBRA scores. Cells with LIBRA score > 0 for any of the antigens used for baiting were defined as SARS-CoV-2-specific.

### Statistical Analysis

The number of samples and subjects used in each experiment are indicated in the figure legends as are the statistical tests used. All tests were performed two-sided. In general, non-parametric Kruskal-Wallis tests were used to test for differences between continuous variables in more than 2 groups and p-values were adjusted for multiple testing using the Dunn’s method. Statistical analysis was performed with Graph-Pad Prism (Version 9.4.1, GraphPad Software, La Jolla California USA) and R (Version 4.1.0). Statistical significance was established at *P* < 0.05.

## ACKNOWLEDGMENTS

We thank the patients, Sara Hasler for assistance with patient recruitment, Laura Bürgi and Rebecca Masek for help with sample processing, the Departments of Otorhinolaryngology and Anesthesiology and the Transplantation Immunology Laboratory of University Hospital Zurich, Esther Baechli, Alain Rudiger, Melina Stüssi-Helbling, Lars C. Huber, the Functional Genomics Center Zurich, the Genomics Facility Basel, Stéphane Chevrier, and Daniel Pinschewer, Klaus Warnatz for helpful discussions and reading of the manuscript, and the members of the Boyman and Moor Laboratories for helpful discussions. Graphical representations were generated with BioRender.com.

## AUTHOR CONTRIBUTION

Y.Z. designed and performed flow cytometry and scRNA-seq experiments, analyzed and interpreted data. J.M. designed and performed scRNA-seq experiments, analyzed and interpreted data. P.T. and S.A. contributed to flow cytometry experiments, patient recruitment and data collection. C.C. contributed to patient recruitment and data collection. M.E.R. and M.S. contributed to patient recruitment and clinical management. I.E.A. analyzed scRNA-seq data. J.N. contributed to patient recruitment. A.E.M. Designed experiments and interpreted data. O.B. conceived the project, designed experiments, and interpreted data. Y.Z. and O.B. wrote the manuscript with contribution by J.M. and A.E.M. All authors edited and approved the final manuscript.

---

**Supplementary Figure 1.**
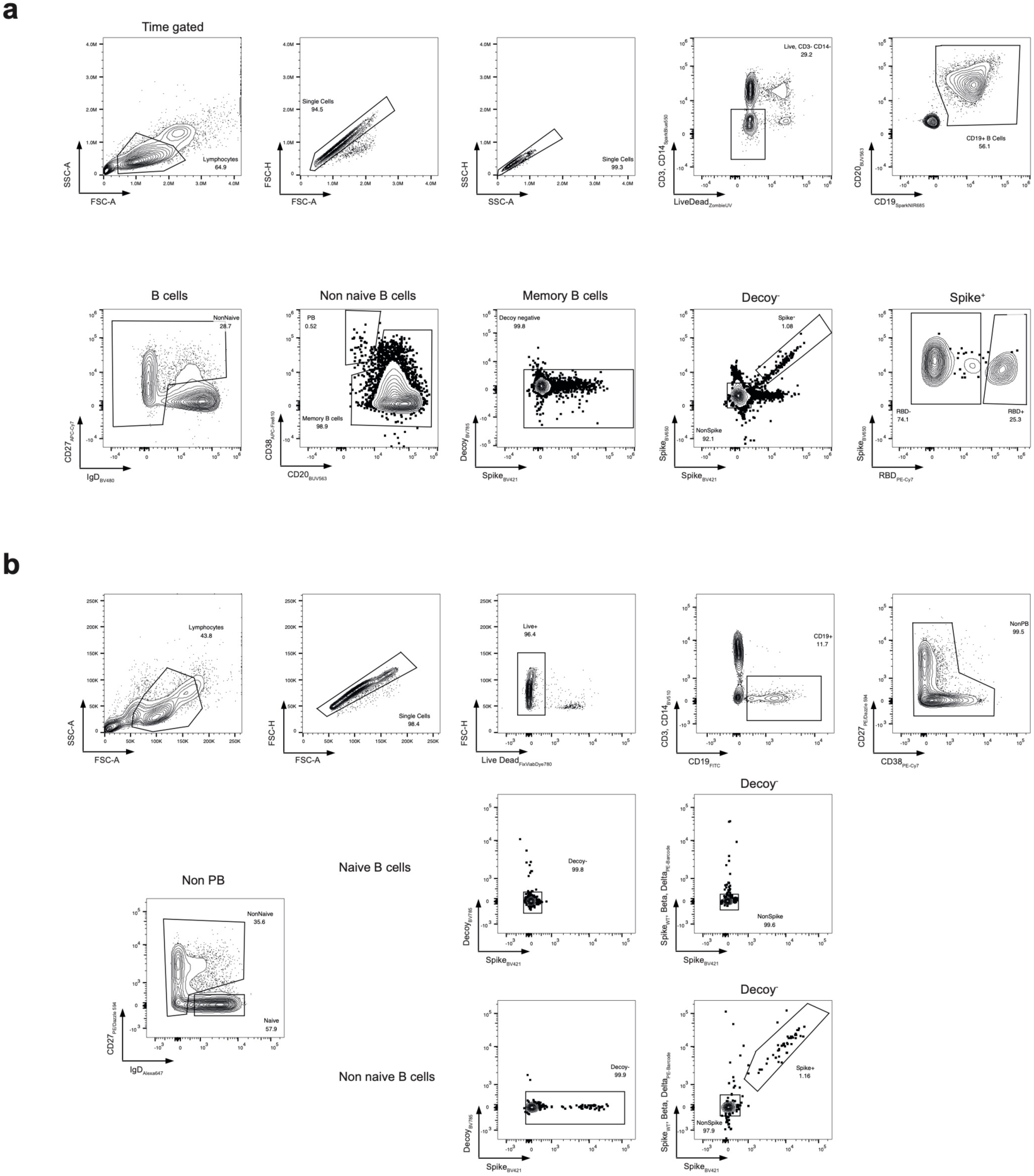
Flow cytometry gating strategies for SARS-CoV-2 spike-specific MBCs. **a**, Gating strategy for SARS-CoV-2 spike^+^ and RBD^+^ memory B cells. **b**, Sorting strategy for SARS-CoV-2 specific and non-specific memory B cells as well as naïve B cell

**Supplementary Figure 2.**
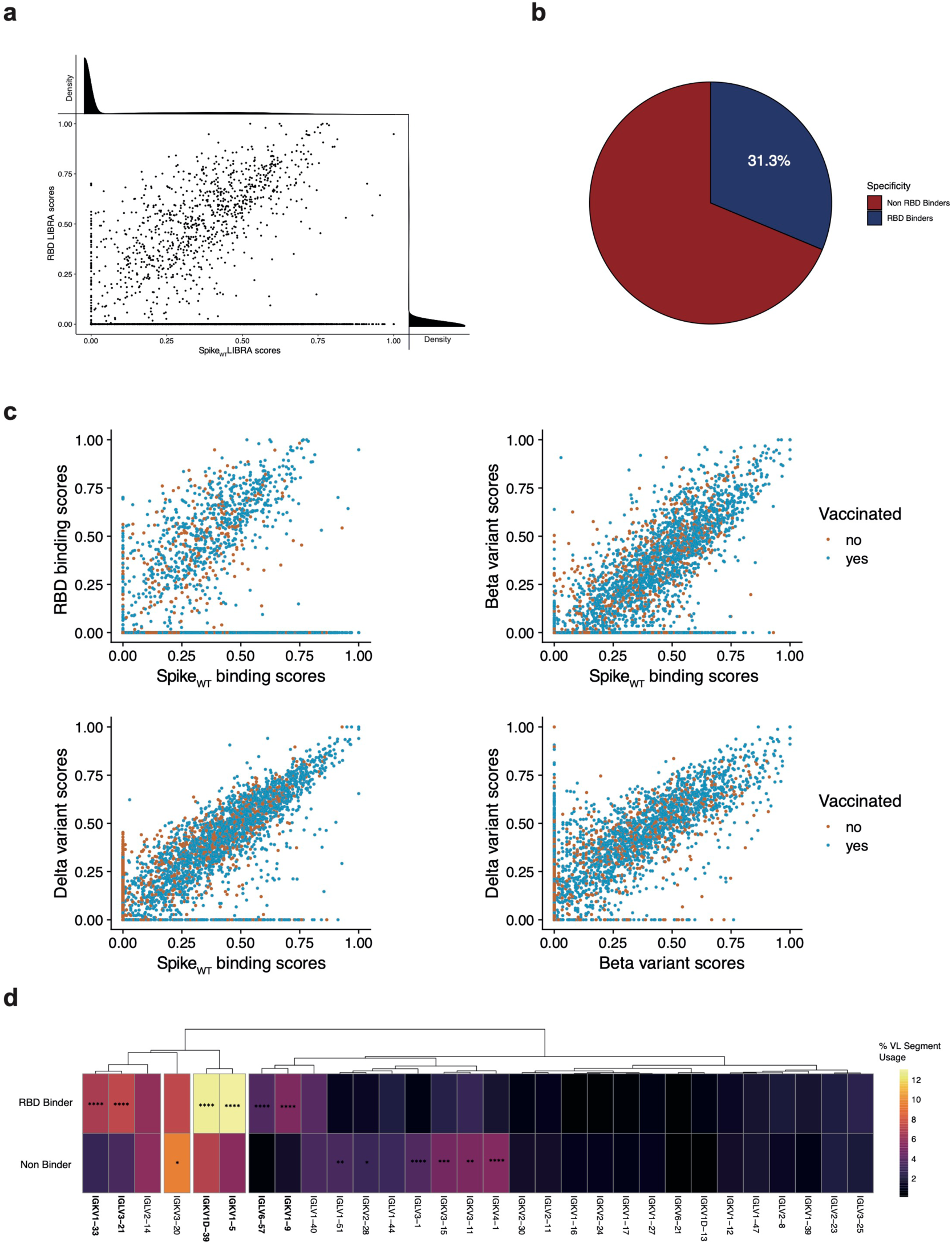
Identification of SARS-CoV-2 spike, RBD and spike variant specific MBCs using scRNA-seq. **a**, Scatter plot comparing LIBRA scores for spike_WT_ and RBD, where every dot represents a cell. Density plots indicating count distributions across LIBRA score ranges are shown for both antigens. **b**, Pie chart showing the percentage of spike_WT_ binders which also bind RBD in the scRNA-seq dataset. **c**, Scatter plots as in **a** showing LIBRA scores for indicated antigen baiting constructs against each other. **d**, Heatmap comparing the V light (VL) gene usage between RBD binders and non-binders. VL segments are sorted by a hierarchical clustering. Colors indicate the frequency within the RBD binders resp. non-binders. The 30 most frequently used segments among RBD binders are shown In **d** frequencies were compared using a two-proportions z-test with a Bonferroni based multiple testing correction. P-values are shown if significant (p<0.05). *p<0.05, **p<0.01, ***p<0.001, **** p<0.0001.

**Supplementary Figure 3.**
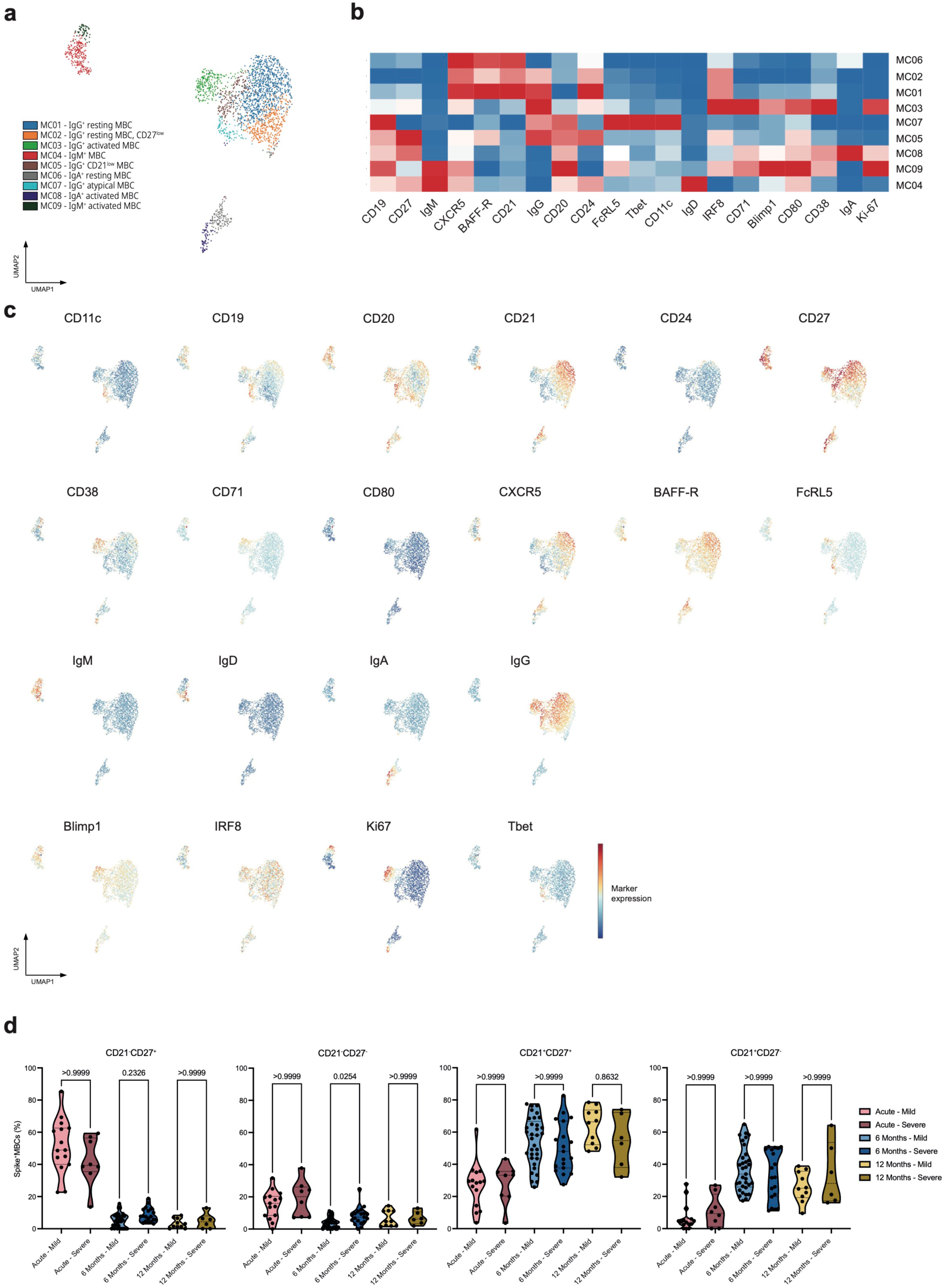
Unsupervised analysis of circulating MBCs after SARS-CoV-2 infection. **a**, UMAP plot of spike^+^ MBCs of all samples which were not vaccinated (n=120), subsampled to maximally 25 cells per sample and colored by clusters identified with a PhenoGraph algorithm. Clusters were manually annotated. **b**, Heatmap of the normalized marker expression from the PhenoGraph clusters. **c**, UMAP as in **a** colored by the indicated marker expression. **d**, Violin plots comparing frequencies of CD21^-^ CD27^+^, CD21^-^CD27^-^, CD21^+^CD27^+^ and CD21^+^CD27^-^ subsets in spike^+^ MBCs separated by disease severity and time points after infection. Mild (acute n=15, six months n=33, 12 months n=10) and severe COVID-19 (acute n=8, six months n=19, 12 months n=6) were compared between the same time point using a Kruskal-Wallis test with a Dunn’s multiple comparison correction, adjusted p-values are shown.

**Supplementary Figure 4.**
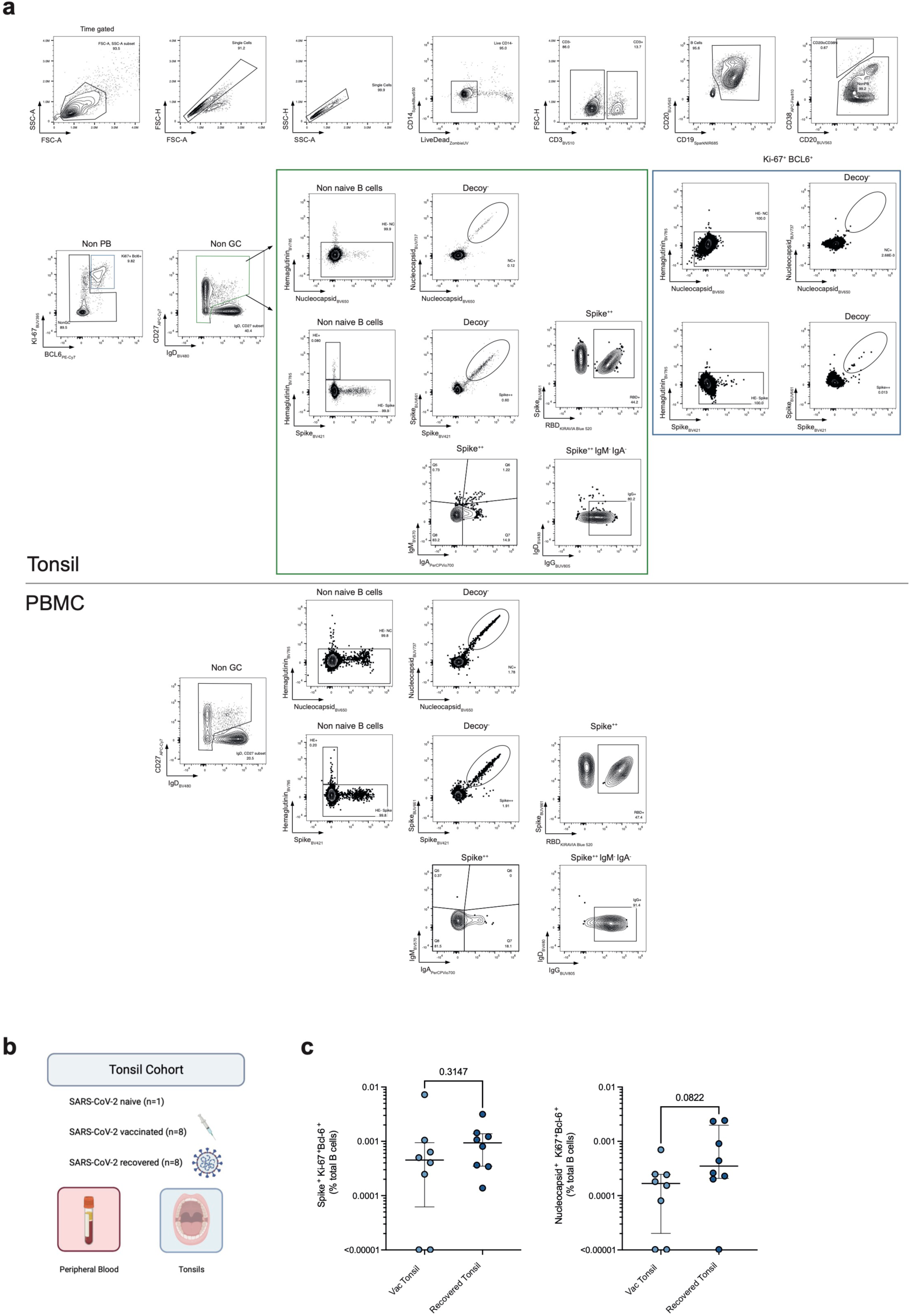
Gating strategy and analysis of tonsillar and circulating B cells. **a**, Gating strategy for the identification of SARS-CoV-2 spike and nucleocapsid-specific germinal center and memory B cells. Shown is the tonsil (top) and the paired peripheral blood sample (bottom) from a COVID-19 recovered individual (CoV-Tissue-02). **b**, Tonsil study cohort overview. **c,** Frequency of spike^+^ (left) and nucleocapsid^+^ (right) germinal center B cells of total B cells in tonsils of SARS-CoV-2 vaccinated and COVID-19 recovered individuals. Frequencies were compared using a Mann Whitney test.

**Supplementary Figure 5.**
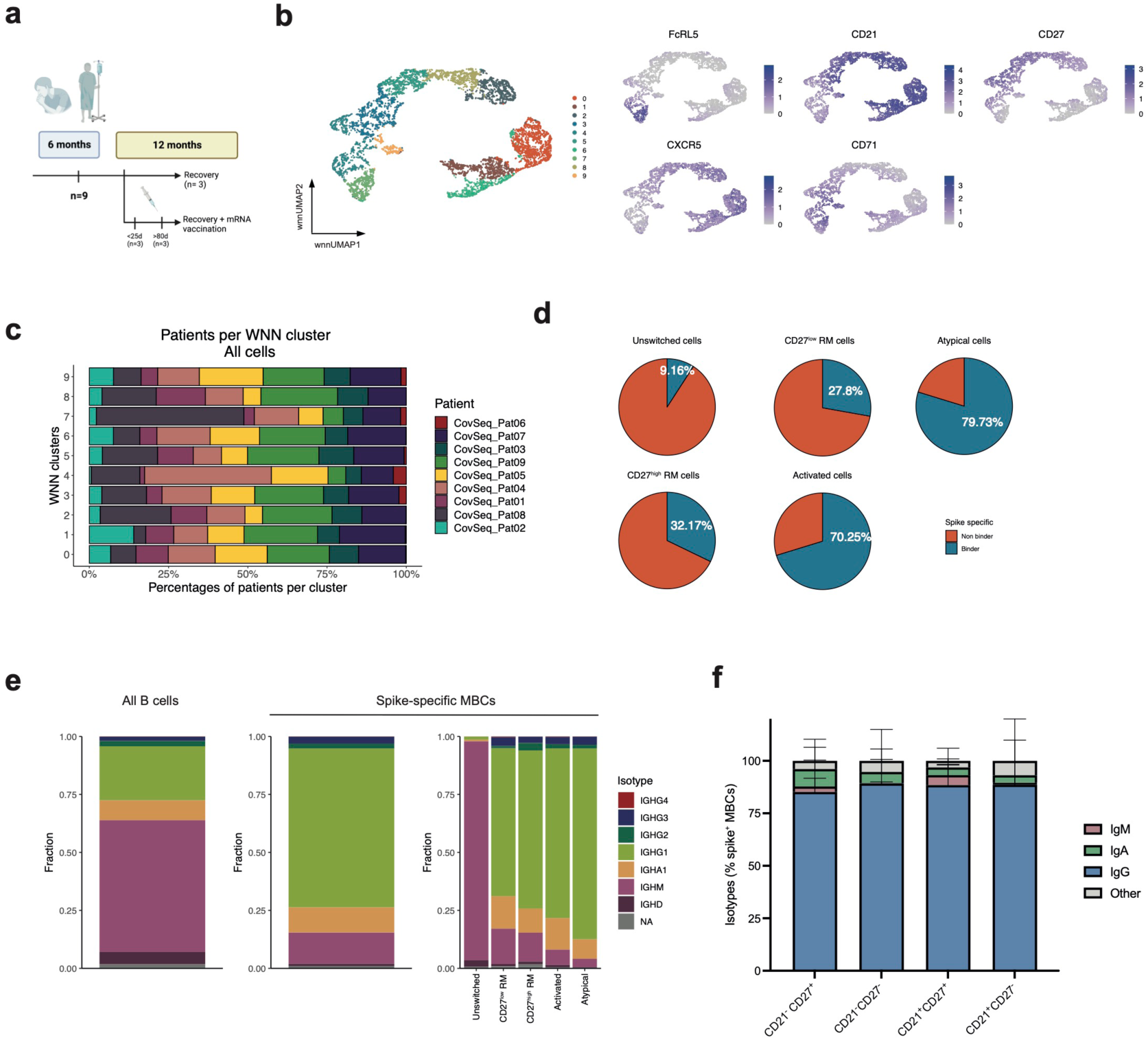
SARS-CoV-2-specific MBC subset identification by scRNA-seq analysis. **a**, scRNA-seq subcohort overview. **b**, Weighted-nearest neighbor UMAP (wnnUMAP) of MBCs from COVID-19 patients at six and at 12 months after infection, colored by clustering based on single-cell transcriptome and cell surface protein information (left) and UMAP colored by indicated surface protein markers (right). **c**, Stacked bar graph showing the single patient contribution to the wnn clusters. **d**, Pie charts showing the percentages of spike^+^ MBCs among all cells in the dataset, separated by MBC subset. **e**, Stacked bar graph showing the isotype and subtype usage in the scRNA-seq dataset on all B cells (left), all spike^+^ (middle) and spike^+^ MBC subsets (right). **f**, Stacked bar graph showing the isotype usage in the spike^+^ MBC subset from the flow cytometry dataset (n=41).

**Supplementary Figure 6.**
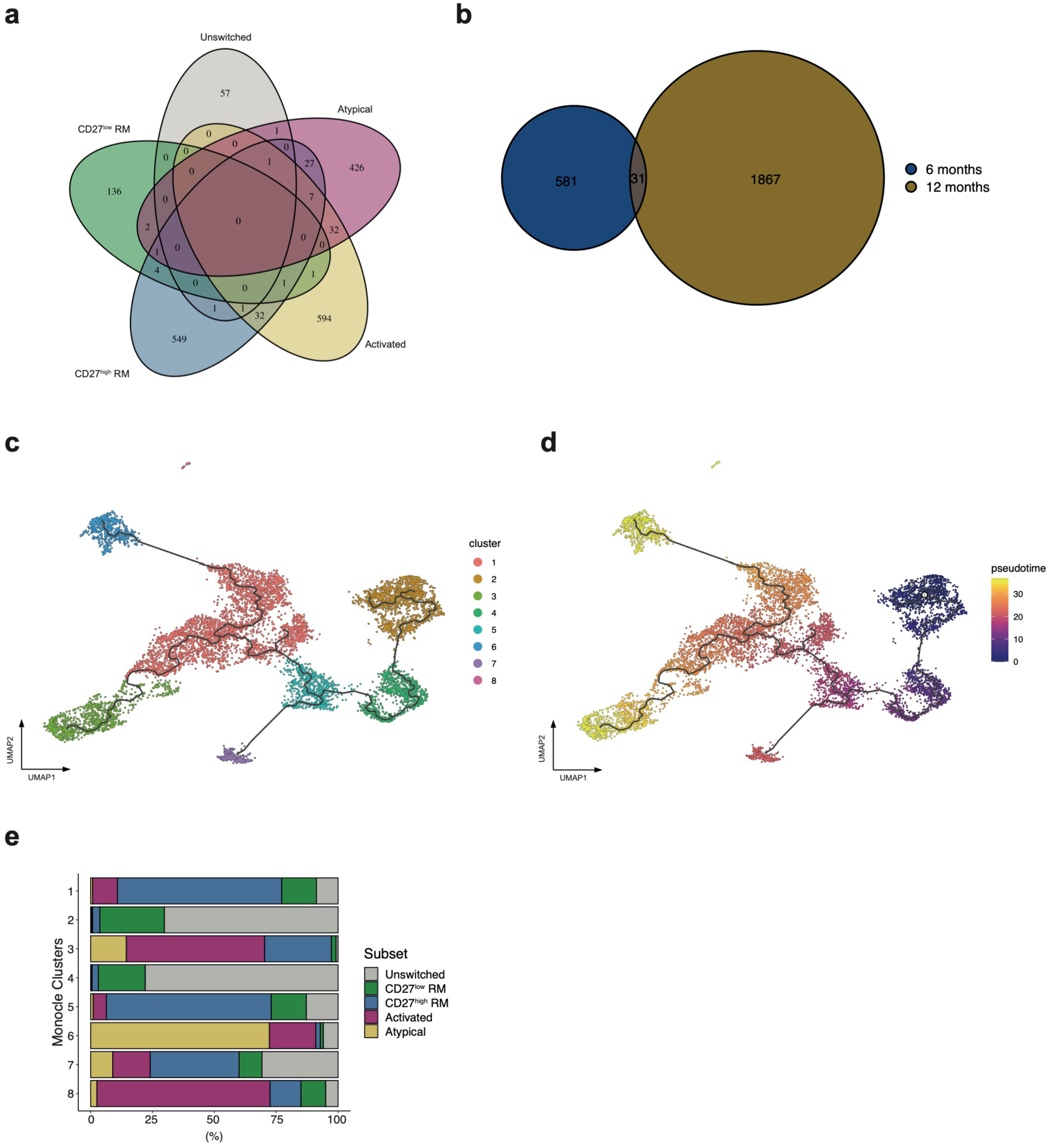
scRNA-seq clonal analysis and Monocle analysis. **a**, Venn diagram showing the clonal overlap of SARS-CoV-2 specific clones in the different MBC subsets. **B**, Venn diagram showing the clonal overlap of SARS-CoV-2 specific clones 6 and 12 months after SARS-CoV-2 infection **c**, UMAP representation of a Monocle analysis on all memory B cells colored by clusters identified via the Monocle algorithm. **d**, UMAP as in **c** colored by a pseudotime annotation. The beginning of the pseudotime was manually set inside the partition with mostly unswitched cells. **e**, Stacked bar graph showing the contribution of SARS-CoV-2 specific MBC subsets to the clusters derived from Monocle.

