## Supplementary Tables for "Fate and plasticity of SARS-CoV-2-specific B cells during memory and recall response in humans"

### Supplementary Table 1. Characteristics of patient cohort.

| **Time point of sampling** | | **Acute infection (n = 59)** | **6 months after acute infection (n = 64)** | | **12 months after acute infection (n = 55)** | |
|---|---|---|---|---|---|---|
| | | | Not vaccinated (n = 61) | Vaccinated (n = 3) | Not Vaccinated (n = 17) | Vaccinated (n = 38) |
| Age, y median (IQR) | | 39 (32 – 58) | 41 (32.5 – 59.5) | 68 (22 – 69) | 39 (30.5 – 60) | 39.5 (32 – 56.5) |
| Sex, female/male | | 30/29 | 30/31 | 3/0 | 7/10 | 18/20 |
| **Severities** | | | | | | |
| Mild | Mild illness, no. | 37 | 36 | 2 | 11 | 27 |
| | Pneumonia, no. | 3 | 3 | 1 | – | 3 |
| Severe | Severe pneumonia, no. | 6 | 9 | – | 2 | 4 |
| | Mild ARDS, no. | 1 | 2 | – | 1 | 1 |
| | Moderate ARDS, no. | 5 | 5 | – | 2 | 1 |
| | Severe ARDS, no. | 7 | 6 | – | 1 | 2 |
| Days after symptom onset, mean (±SD) | | 13.71 (±9.51) | 201.9 (±28.94) | 208.7 (±29.67) | 374.2 (±23.44) | 377.6 (±22.5) |
| **SARS-CoV-2 Vaccinations**[a] | | | | | | |
| Moderna, no. (1/2 doses) | | – | – | 1/0 | – | 20/3 |
| BioNTech, no. (1/2 doses) | | – | – | 0/2 | – | 7/7 |
| Days after vaccination, mean (±SD) | | – | – | 16 (±6.25) | – | 72.82 (±48.35) |
| **SARS-CoV-2 Antibodies** | | | | | | |
| Spike S1 IgA, OD ratio mean (±SD) | | 4.53 (±4.02) | 3.65 (±2.51) | 8.36 (±0.38) | 3.45 (±2.91) | 8.97 (±1.45) |
| Spike S1 IgG, OD ratio mean (±SD) | | 3.09 (±3.77) | 4.28 (±3.01) | 11.24 (±0.68) | 4.0 (±3.48) | 10.0 (±1.22) |

Abbreviations: IQR, interquartile range; OD, optical density; SD, standard deviation.
[a] For one patient it was not retrievable which SARS-CoV-2 mRNA vaccine the patient received.

**Supplementary Table 2. Characteristics of subcohort used for single-cell RNA sequencing.**

| Patient ID | Sex | Timepoint | Age [y] | COVID-19 Severity | Days after symptom onset | Vaccination | Days after vaccination |
|---|---|---|---|---|---|---|---|
| CovSeq_Pat01 | Female | 6 months | 59 | Mild | 214 | Not vaccinated | – |
| | | 12 months | 60 | Mild | 408 | Not vaccinated | – |
| CovSeq_Pat02 | Female | 6 months | 69 | Sev ARDS | 117 | Not vaccinated | – |
| | | 12 months | 70 | Sev ARDS | 312 | Not vaccinated | – |
| CovSeq_Pat03 | Male | 6 months | 59 | Mod ARDS | 255 | Not vaccinated | – |
| | | 12 months | 59 | Mod ARDS | 394 | Not vaccinated | – |
| CovSeq_Pat04 | Female | 6 months | 42 | Mild | 181 | Not vaccinated | – |
| | | 12 months | 42 | Mild | 377 | Moderna, 1 dose | 9 |
| CovSeq_Pat05 | Male | 6 months | 53 | Mild | 177 | Not vaccinated | – |
| | | 12 months | 53 | Mild | 372 | Moderna, 1 dose | 11 |
| CovSeq_Pat06 | Male | 6 months | 76 | Sev Pneumonia | 203 | Not vaccinated | – |
| | | 12 months | 77 | Sev Pneumonia | 384 | Moderna, 1 dose | 23 |
| CovSeq_Pat07 | Female | 6 months | 32 | Mild | 189 | Not vaccinated | – |
| | | 12 months | 32 | Mild | 364 | Moderna, 1 dose | 85 |
| CovSeq_Pat08 | Female | 6 months | 54 | Mild | 282 | Not vaccinated | – |
| | | 12 months | 54 | Mild | 415 | Moderna, 1 dose | 87 |
| CovSeq_Pat09 | Male | 6 months | 27 | Mild | 193 | Not vaccinated | – |
| | | 12 months | 27 | Mild | 377 | Moderna, 1 dose | 108 |

### Supplementary Table 3. Characteristics of cohort used for analysis of tonsillar immune cells.

| | **Vaccinated (n = 8)** | **Recovered (n = 8)** | **Naïve (n = 1)** |
|---|---|---|---|
| Age, y median (IQR) | 33 (25.5 – 43.5) | 25 (22 – 37) | 65 |
| Sex, female/male | 2/6 | 7/1 | 0/1 |
| Anamnestic COVID-19, no. | – | 6 | – |
| **SARS-CoV-2 Vaccinations** | | | |
| Vaccine doses, no. (0/1/2/3) | 0/0/6/2 | 2/1/2/3 | – |
| Vaccine, no. (Moderna/ BioNTech /Mixed mRNA/Unkown) | 6/0/0/2 | 1/4/1/0 | – |
| Days after last vaccination, mean (±SD) | 144.1 (±51.61) | 117.5 (±56.53) | – |
| **Anti-SARS-CoV-2 Antibodies** | | | |
| Anti-S1 IgG (SOC), mean (±SD) | 131.5 (±94.54) | 177.9 (±98.08) | 0.37 |
| Anti-S2 IgG (SOC), mean (±SD) | 2.4 (±1.21) | 5.17 (±2.76) | 0.47 |
| Anti-RBD IgG (SOC), mean (±SD) | 107.8 (±62.51) | 134.0 (±73.29) | 0.03 |
| Anti-NC IgG (SOC), mean (±SD) | 0.16 (±0.03) | 22.41 (±29.12) | 0.12 |
| Abbreviations: IQR, interquartile range; SD, standard deviation; SOC, signal-over cut-off. | | | |

**Supplementary Table 4. Antibodies used for B cell staining.**

| Antigen | Fluorophore | Provider | Cat No |
|---|---|---|---|
| CD3 | SparkBlue550 | Biolegend | 344852 |
| CD11c | BUV615 | BD Biosciences | 612967 |
| CD14 | SparkBlue550 | Biolegend | 367147 |
| CD19 | SparkNIR 685 | Biolegend | 302270 |
| CD20 | BUV563 | BD Biosciences | 748456 |
| CD21 | BUV496 | BD Biosciences | 750614 |
| CD24 | BUV805 | BD Biosciences | 742010 |
| CD27 | APC-Cy7 | Biolegend | 356424 |
| CD38 | APC-Fire810 | Biolegend | 303549 |
| CD71 | PerCP-Cy5.5 | Biolegend | 334114 |
| CD80 | PE-Cy5 | Biolegend | 305209 |
| CXCR5 | BV750 | Biolegend | 356941 |
| BAFF-R | BV605 | BD Biosciences | 743571 |
| FcRL5 | BUV615 | BD Biosciences | 751131 |
| IgD | BV480 | BD Biosciences | 566187 |
| IgM | BV570 | Biolegend | 314517 |
| IgA | APC | Miltenyi Biotec | 130-113-472 |
| IgG | BUV737 | BD Biosciences | 741858 |
| IgG1 | PE | Cytognos | CYT-IGG1PE |
| IgG3 | FITC | Cytognos | CYT-IGG3F |
| BLIMP1 | PE-Dazzle594 | BD Biosciences | 565274 |
| IRF8 | V450 | Invitrogen | 48-9852-82 |
| Ki67 | BUV395 | BD Biosciences | 564071 |
| Tbet | BV711 | Biolegend | 644819 |
| ZombieUV | | Biolegend | 423107 |
| Streptavidin | BV421 | Biolegend | 405226 |
| Streptavidin | BV650 | Biolegend | 405231 |
| Streptavidin | BV785 | Biolegend | 405249 |
| Streptavidin | PE-Cy7 | Biolegend | 405206 |

**Supplementary Table 5. Antibodies used for fluorescently-activated cell sorting.**

| Antigen | Fluorophore | Provider | Cat No |
|---|---|---|---|
| CD3 | BV510 | Biolegend | 317332 |
| CD14 | BV510 | Biolegend | 301841 |
| CD19 | FITC | Biolegend | 302206 |
| CD21 | TotalSeq™-C0181 | Biolegend | 354923 |
| CD27 | PE-Dazzle | Biolegend | 356421 |
| CD27 | TotalSeq™-C0154 | Biolegend | 302853 |
| CD38 | PE-Cy7 | Biolegend | 303515 |
| IgD | Alexa647 | Biolegend | 348227 |
| IgD | TotalSeq™-C0384 | Biolegend | 348245 |
| CD71 | TotalSeq™-C0394 | Biolegend | 334125 |
| CXCR5 | TotalSeq™-C0144 | Biolegend | 356939 |
| FcRL5 | TotalSeq™-C0829 | Biolegend | 340309 |
| Hashtag 1 | TotalSeq™-C0251 | Biolegend | 394661 |
| Hashtag 2 | TotalSeq™-C0252 | Biolegend | 394663 |
| Hashtag 3 | TotalSeq™-C0253 | Biolegend | 394665 |
| Hashtag 4 | TotalSeq™-C0254 | Biolegend | 394667 |
| Hashtag 5 | TotalSeq™-C0255 | Biolegend | 394669 |
| Hashtag 6 | TotalSeq™-C0256 | Biolegend | 394671 |
| Hashtag 7 | TotalSeq™-C0257 | Biolegend | 394673 |
| Hashtag 8 | TotalSeq™-C0258 | Biolegend | 394675 |
| Hashtag 9 | TotalSeq™-C0259 | Biolegend | 394677 |
| Hashtag 10 | TotalSeq™-C0260 | Biolegend | 394679 |
| Streptavidin | TotalSeq™-C0951 PE | Biolegend | 405261 |
| Streptavidin | TotalSeq™-C0952 PE | Biolegend | 405263 |
| Streptavidin | TotalSeq™-C0954 PE | Biolegend | 405267 |
| Streptavidin | TotalSeq™-C0971 | Biolegend | 405271 |
| Streptavidin | TotalSeq™-C0972 | Biolegend | 405273 |
| Streptavidin | BV421 | Biolegend | 405226 |
| Streptavidin | BV785 | Biolegend | 405249 |
| Fixable Viability Dye | eFluor™ 780 | Invitrogen | 65-0865-14 |

**Supplementary Table 6. Antibodies used for staining of tonsillar samples.**

| Antigen | Fluorophore | Provider | Cat No |
|---|---|---|---|
| CD3 | BV510 | Biolegend | 317332 |
| CD14 | SparkBlue550 | Biolegend | 367147 |
| CD19 | SparkNIR 685 | Biolegend | 302270 |
| CD20 | BUV563 | BD Biosciences | 748456 |
| CD21 | BUV496 | BD Biosciences | 750614 |
| CD27 | APC-Cy7 | Biolegend | 356424 |
| CD38 | APC-Fire810 | Biolegend | 303549 |
| CD45RB | APC | Invitrogen | MA1-19461 |
| CD69 | BV605 | Biolegend | 310937 |
| CXCR4 | PE-Cy5 | Biolegend | 306507 |
| CXCR5 | BV750 | Biolegend | 356941 |
| FcRL5 | BUV615 | BD Biosciences | 751131 |
| IgD | BV480 | BD Biosciences | 566187 |
| IgM | BV570 | Biolegend | 314517 |
| IgG | BUV805 | BD Biosciences | 742041 |
| IgG1 | PE | Cytognos | CYT-IGG1PE |
| IgA | PerCP-Vio700 | Miltenyi Biotec | 130-114-004 |
| BCL6 | PE-Cy7 | Biolegend | 358511 |
| BLIMP1 | PE-Dazzle594 | BD Biosciences | 565274 |
| Ki67 | BUV395 | BD Biosciences | 564071 |
| Tbet | BV711 | Biolegend | 644819 |
| ZombieUV | | Biolegend | 423107 |
| Streptavidin | BUV661 | BD Biosciences | 612979 |
| Streptavidin | BUV737 | BD Biosciences | 612775 |
| Streptavidin | BV421 | Biolegend | 405226 |
| Streptavidin | BV650 | Biolegend | 405231 |
| Streptavidin | BV785 | Biolegend | 405249 |
| Streptavidin | KIRAVIA Blue 520 | Biolegend | 405171 |
